# From strandline vegetation to embryo dunes: natural sand retention along a heavily urbanized coast

**DOI:** 10.64898/2026.07.14.738448

**Authors:** Charlotte Taelman, Sam Provoost, Femke Batsleer, Jan Van Uytvanck, Dries Bonte

**Author notes:** Corresponding author: C. Taelman, Ghent University, Department of Biology, Terrestrial Ecology Unit, K. L. Ledeganckstraat 35, 9000 Ghent, Belgium. These 2 two authors share first-authorship (Charlotte Taelman, Sam Provoost).

## Abstract

1. Sandy beaches along urbanized coasts are increasingly managed through beach nourishment and hard infrastructure, yet these interventions often constrain natural dune-building processes. Along the Belgian coast, where much of the beach–dune interface is bordered by dikes, promenades and intensive recreation, strandline vegetation may provide an overlooked mechanism for retaining sand and initiating embryo dune development.
2. We assessed the potential for four pioneer dune plant species (*Cakile maritima*, *Calamagrostis arenaria*, *Elymus farctus* and *Salsola kali*) to establish, develop vegetation cover and contribute to sand accumulation along the Belgian coast. Using field surveys from 2017–2023, LiDAR-derived beach elevation and annual sediment dynamics, we modelled species occurrence and abundance/cover in low-disturbance reference zones and projected these relationships across the wider coastline.
3. Occurrence models identified where abiotic conditions allow plants to establish and persist until the late growing season, whereas zero-inflated abundance/cover models estimated expected vegetation development across environmental gradients. Predicted occurrence was widespread for several species, suggesting that the abiotic gradients modelled here are not the primary constraints on potential establishment across large parts of the coast. In contrast, expected abundance/cover showed stronger species-specific responses, particularly to sand accretion, indicating that sediment dynamics mainly affect post-establishment vegetation development rather than occurrence alone.
4. Independent field measurements of embryo dunes showed positive relationships between vegetation cover and local sand accumulation for all four species. When scaled using spatial predictions of potential abundance/cover, pioneer vegetation could retain substantial volumes of sand, with *Cakile maritima* contributing the largest share, followed by *Salsola kali*, *Elymus farctus* and *Calamagrostis arenaria*. Estimated volumes depended on assumptions about whether vegetation occurs as dispersed units or aggregated patches.
5. **Synthesis and applications.** Our results show that, even along a heavily urbanized and nourished coastline, abiotic conditions can support strandline vegetation and embryo dune initiation where disturbance is reduced. Management actions such as limiting trampling, adapting beach cleaning and protecting strandline vegetation could enhance the retention of nourished sand and support nature-based coastal defense. Rather than replacing engineered interventions, strandline vegetation may increase the efficiency with which available sediment is retained within the beach–dune system.

## Introduction

Sandy beaches are dynamic coastal ecosystems that cover one third of all ice-free shorelines worldwide (Luijendijk et al., 2018). Their morphology is shaped by the interaction of wind, waves, tides and sea-level fluctuations (Athanasiou et al., 2020), creating environmental gradients that support specialized biotic communities (Defeo et al., 2009). Globally, sandy shorelines provide substantial ecological and socio-economic value: they host distinct biodiversity, support local economies through beach tourism (Klein et al., 2004) and act as natural buffers protecting coastal communities from storm surge flooding (Athanasiou et al., 2020).

Despite their importance, sandy beach ecosystems are increasingly subjected to pressures operating at multiple spatial and temporal scales. Coastal armoring (e.g., dikes and promenades) (Dugan et al., 2008), beach nourishment to compensate for sand losses (de Schipper et al., 2020; Speybroeck et al., 2006), non-selective beach cleaning and infrastructure development (Defeo et al., 2009; Lansu et al., 2024; Nordstrom, 2000) alter sediment dynamics and reduce the habitat quality. In addition, global, climate-driven stressors such as sea level rise and increased storms (Athanasiou et al., 2020; Paprotny et al., 2021; Vousdoukas et al., 2020) further enhance erosion shorelines worldwide (Feagin et al., 2005).

Sandy beaches form a dynamic interface between marine and terrestrial (eco-)systems through the cross-shore transport and retention of sediment and nutrients (Defeo et al., 2009; Short & Jackson, 2013). Despite the harsh environmental conditions, pioneer vascular plant species can establish on the dry beach and add a determining, biotic element to the system. Their establishment is facilitated through the availability of (freshwater) soil moisture (Homberger et al., 2025) and nutrients derived from decay of deposited marine wrack (van Egmond et al., 2019), with nitrogen playing a central role physiological processes underlying salt tolerance (Pakeman & Lee, 1991). Conversely, prolonged marine inundation and excess sediment dynamics (sedimentation or erosion) limit the establishment and survival of vascular plants (Bonte et al., 2021).

Once established, pioneer plants initiate early dune development by trapping wind-blown sand, and forming small shadow dunes (Hesp & Smyth, 2017; Maun, 2009). Through this bio-geomorphic feedback, vegetation enhances local sand accumulation, leading to the formation of embryo dunes. The subsequent development of these dunes follows a seasonal cycle, driven by sand supply and vegetation growth in summer, and erosion and plant decline during winter (Montreuil et al., 2013; van Puijenbroek et al., 2017). Ultimately, the interaction between vegetation, marine wrack input, and hydrodynamic and aeolian processes shapes beach morphology and influences sediment retention at the land–sea interface (Silva et al., 2016).

The Belgian coast is among the most intensively used sandy shorelines in Europe, with a long tradition of coastal urbanization, including the construction of dikes, promenades, and large housing complexes since the 19^th^ century (Pillen et al., 2017). As a result, approximately 60 % of the beach-dune interface is now occupied by human infrastructure. These modifications strongly constrain opportunities for natural embryo dune formation. Mechanical beach cleaning, removal of wrack and vegetation, artificial beach reprofiling, temporary recreational infrastructure and intensive trampling all reduce the establishment and persistence of pioneer vegetation (Speybroeck & Bonte, 2008).

At the same time, accelerating sea-level rise and increasing storm frequency are enhancing erosion pressure on sandy coasts worldwide, driving the need for effective coastal protection strategies. Along the Belgian coast, this is primarily achieved through a combination of hard infrastructure (e.g. dikes) and soft engineering approaches such as beach nourishments, which are used to compensate for structural sediment losses (MDK, 2011). Over the past decades, nourished sand volumes have been substantial, averaging approximately 500.000 m³ per intervention, with considerable variation among sites and years (Toon Verwaest – Flanders hydraulics, personal communications). While nourishments maintain sediment availability within the coastal system, the extent to which this sediment is retained on the upper beach, or redistributed by marine and aeolian processes, remains uncertain. In this context, strandline vegetation may play an important role by trapping wind-blown sand and enhancing local sediment retention, thereby potentially reinforcing natural dune-building processes. In parallel, policy developments such as the EU Nature Restoration Law (Sundseth, 2025) emphasize the need to restore degraded coastal habitats. Achieving these objectives requires a robust understanding of the environmental conditions that allow pioneer vegetation to establish and initiate dune development along heavily urbanized coasts.

In this paper, we investigate the beaches’ physical boundary conditions governing the establishment and persistence of several pioneer plant species. Using occurrence and abundance/cover data from low-disturbance environments, we model their ecological niches based on key abiotic drivers and project these conditions across the entire coast to estimate potential habitat suitability under minimal human interference. In addition, we quantify the relationship between vegetation cover and sand accumulation to assess the potential contribution of embryo dune formation to coastal sediment retention. Rather than replacing engineered interventions, we evaluate how these natural processes may complement existing coastal management strategies by enhancing the retention of available sediment.

## Materials and methods

### Niche model

To investigate the environmental conditions governing strandline plant establishment, we conducted field surveys to collect data on dry beach plant species along the entire coastline of Belgium. This coastline consists of sandy beaches backed by coastal dunes, interspersed with extensive urban infrastructure, including dikes, promenades, harbors and built-up areas. Surveys were carried out annually between 2017 and 2023 during the late growing season (August-November), thereby capturing realized plant abundance following establishment rather than initial germination.

The study focused on the dry beach zone, defined as the zone between the springtide high waterline (named SHW hereafter) and the marram foredune toe (Fig. 1, top). Within this zone, we recorded occurrences of the four most common pioneer species (Fig. 1, bottom): sea rocket (*Cakile maritima*), marram grass (*Calamagrostis arenaria*), sand couch grass (*Elymus farctus*) and prickly sandwort (*Salsola kali*).

**Fig. 1.**
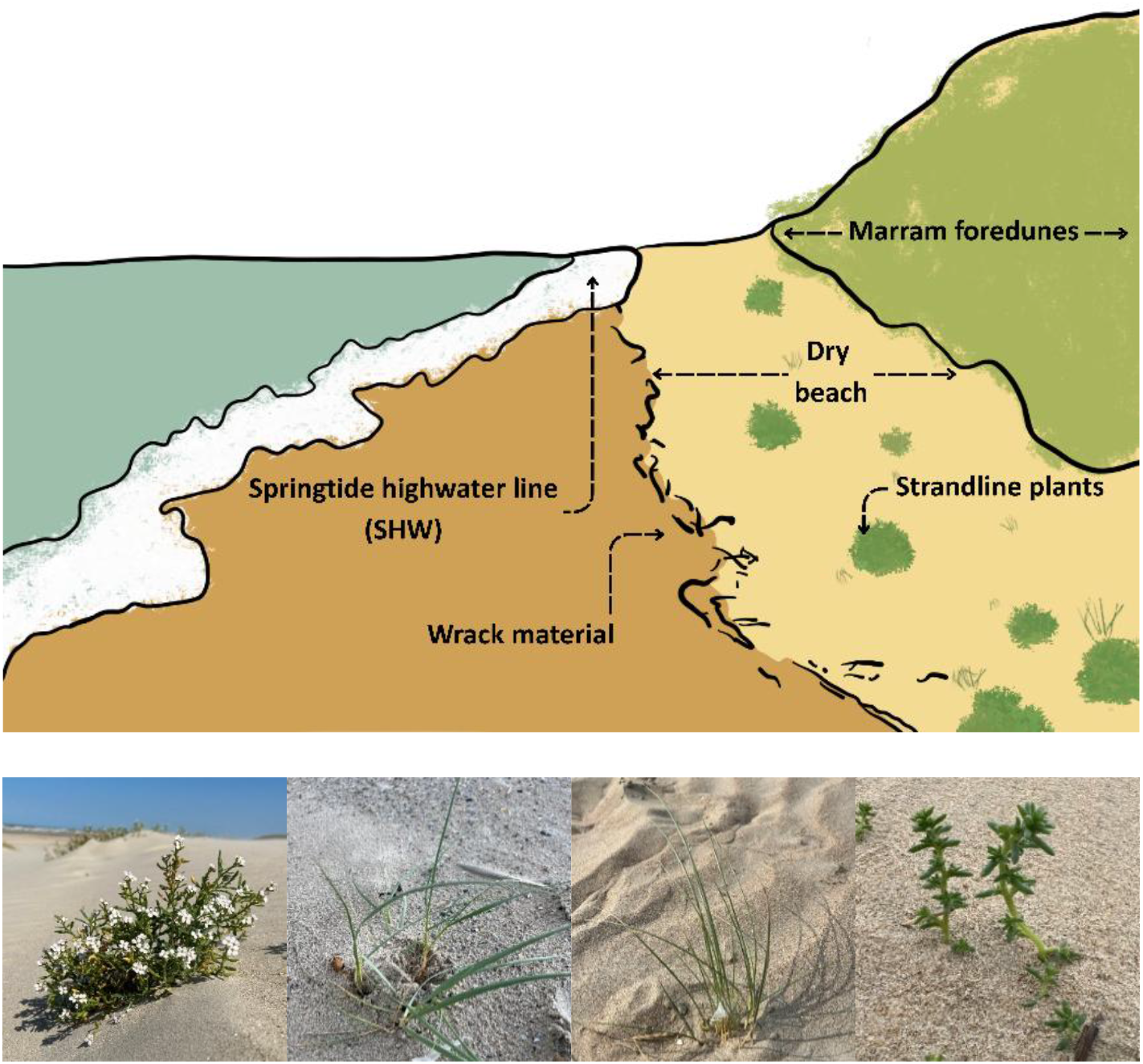
Top: Schematic illustration of the sampled environment on the beach. Plant surveys were conducted in the area indicated as “dry beach”, located between the wrack line and foredune toe. (Drawn by Charlotte Taelman). Bottom: the four species sampled in our survey. From left to right: Cakile maritima, Elymus farctus, Calamagrostis arenaria, and Salsola kali.

Species occurrences were collected as georeferenced point data (EPSG:4326) using the mobile applications iObs (Apple) and ObsMapp (Android) from the citizen-science platform observations.be. Data were collected along the full coastline, from the French border to the Dutch border. Sampling design varied among years: surveys were conducted either as continuous alongshore transects (2017, 2022–2023) or as approximately 17 sub-transects of ∼250 m distributed along the coastline (2018– 2021). All observations were spatially explicit and treated consistently within the modelling framework.

At each observation point, species identity and local abundance/cover were recorded. Field observations consisted of georeferenced presence records only: plants were recorded where they were observed, but true absences were not surveyed directly. Abundance was expressed using categorical classes corresponding to the surface area occupied by the plant patch (S.1.2), allowing standardized comparison among species with contrasting growth forms, including annual species and clonally expanding perennial grasses. These categorical values were subsequently converted to the median value of each class to obtain a numerical, count-like estimate of local abundance/cover. These values should therefore be interpreted as proxies for local plant cover or relative abundance, rather than as true counts of individuals, especially for clonally spreading grasses such as *C. arenaria* and *E. farctus*.

Because the raw field observations contained only presences, additional zero/background locations were required to model species occurrence and abundance across the beach landscape. For the occurrence model, we generated ecologically defined pseudo-absences within a maximum distance of 100 m from observed presences, while excluding the immediate vicinity of each observation using a 5 m buffer (S.1.7). For each species and year, the number of pseudo-absences was matched to the number of observed presences, resulting in a balanced presence–pseudo-absence dataset for modelling occurrence. The occurrence and abundance/cover models used zero/background information differently. The occurrence model explicitly used the balanced presence–pseudo-absence dataset to estimate the probability of presence. The abundance/cover model then used the presence-only dataset, without the added pseudo-absences, where the presence-observations were assigned the median value of the corresponding abundance/cover class (S.1.2).

Spatial data visualizations, processing, and map making of the geolocated data were performed in QGIS (v3.44.5, QGIS.org, 2026. QGIS Geographic Information System. QGIS Association. http://www.qgis.org). We further used R (v4.4.3, R Core Team, 2025) for data cleaning (*tidyverse, (Wickham et al., 2019)),* data analysis (*terra (Hijmans, 2025), sf and sp (Pebesma & Bivand, 2023)*, *R-INLA (Engel et al., 2022; Lindgren et al., 2011)*) and producing graphs (*showtext (Qio, 2024), ggplot2 (Wickham, 2016))*.

Environmental predictors were derived from digital terrain models (DTMs) of the Belgian coast provided by the Flemish government (Department Mobility and Public Services, MDK). These DTMs were generated through airborne LiDAR conducted usually around spring (March-June) and fall-winter (October-January). An overview of the timing of each LiDAR survey can be found in S.1.3.

To quantify exposure to marine inundation, we calculated beach elevation relative to the springtide high waterline (SHW). SHW was estimated using tide measurements from three coastal stations (Nieuwpoort, Oostende, Zeebrugge) over the period 2000 and 2010 (MDK, 2010), and interpolated to a 100-meter resolution raster covering the entire Belgian coast. We assume that spatial patterns in SHW remain stable over time at the scale relevant to this study.

We subtracted the SHW-elevation from the spring DTMs for each year of sampling (*Raster Calculator*, QGIS) to obtain the elevation of the plants on the beach, relative to the SHW (equation 1).

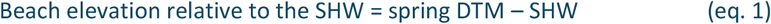

To capture annual sediment dynamics (erosion and/or sand burial), we calculated the difference between fall–winter and spring DTMs (equation 2).

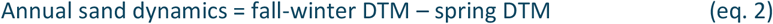

Positive values indicate net sand accumulation (burial), while negative values indicate erosion. All raster layers had a spatial resolution of 1 m, and missing data were not interpolated.

To isolate the effects of physical boundary conditions from human disturbance, we defined 10 regions of interest (ROIs) along the coast, consisting of beach sections at least 1 km in length (except for the bay of Heist) situated in front of well-developed foredune systems. These areas were delineated using high-resolution orthophotos (RGB, 0.15 m). Species observations were restricted to these ROIs, which are characterized by relatively low levels of trampling and mechanical beach cleaning is usually absent. This approach assumes that abiotic constraints on plant establishment can be inferred from these low-disturbance environments and subsequently extrapolated to the more intensively used parts of the coastline.

To quantify environmental conditions at each observation, we extracted mean values of elevation relative to SHW and annual sand dynamics within a 2.5 m buffer around each datapoint. This radius was chosen to reflect the typical spatial extent of individual plants or small patches (∼5 m diameter), and to better represent the local environment experienced by the vegetation. Both environmental variables were standardized (z-scores) prior, to facilitate comparisons of effect size and ensure model stability, including the corresponding quadratic terms.

We used a two-component spatial modelling framework to distinguish between species occurrence and local abundance/cover. First, species-specific occurrence was modelled as a binary response using the presence–pseudo-absence dataset described above. Occurrence was fitted with a binomial model with a logit link (“binomial” in R-INLA), thereby estimating the probability that abiotic conditions allow a species to establish. Second, species-specific local abundance/cover was modelled using the strictly positive class-median values as described above, to estimate abundance responses along environmental gradients. To account for high overdispersion, we fitted a negative-binomial count distribution with a log link. We fitted the model using the “zero-inflated negative binomial (0)” distribution in R-INLA. Although no zero abundances were present in the data, this formulation corrects for overdispersion while still allowing predictions of zeros at unsampled locations. Consequently, the zero-inflation component was not informed by observed zeros in the dataset, but the model specification still allows the prediction of zero abundances in the environmental space.

Both model components included the same abiotic predictors: linear and quadratic effects of annual sand dynamics and elevation relative to the springtide high-water line (SHW) (equation 3-4).

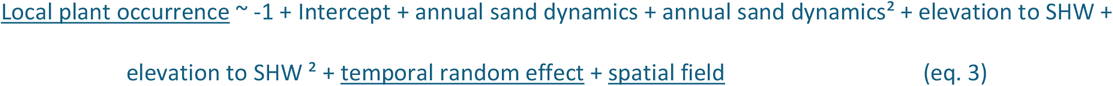

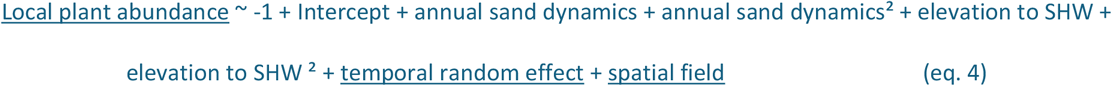

Ecologically, the occurrence model identifies where plants can establish and persist until the late growing season, whereas the abundance/cover model describes how much vegetation is expected to develop across the full beach landscape. Zeros in the abundance model may reflect abiotic unsuitability, but also spatially structured non-abiotic constraints such as disturbance, beach cleaning, trampling, dispersal limitation, or non-detection. Residual spatial structure was accounted for using a Matérn spatial random field implemented through the SPDE approach in R-INLA (Rue et al., 2009). Temporal autocorrelation among sampling years was modelled using a first-order autoregressive effect (AR1). The spatial mesh was constructed using *meshbuilder* (R-INLA, (Engel et al., 2022; Lindgren et al., 2011)) based on the spatial distribution of observations.

Models were fitted separately for each species. Model performance and robustness were assessed using repeated cross-validation (10 iterations), with 70% of the data used for training and 30% for testing in each run. Computations were performed using the high-performance computing infrastructure of Ghent University (HPC-UGent), and posterior distributions, hyperparameters, and spatial fields were retained for further analysis.

### Predictions of the potential occurrence and abundance of plants

We generated both non-spatial and spatially explicit predictions using the fitted INLA models. First, we performed non-spatial predictions to explore species responses across environmental gradients. For each model iteration, we predicted the species occurrence and abundance/cover for a series of standardized values of annual sand dynamics ([-3, 3]) and elevation relative to SHW ([-3, 16]) (n=46101 combinations, step size = 0.05m) covering the observed range of environmental conditions (S.1.8).

Second, we performed spatially explicit predictions across the entire Belgian coastline (including dike-beaches) that consider the modelled spatial fields. These predictions were based on the modeled relationships between species abundance/occurrence and abiotic conditions. This projection assumes that species–environment relationships inferred from low-disturbance ROIs are transferable to the wider coastline. To define the spatial extent of predictions, we extracted the Belgian Coastal Zone from the Corine Landcover classification (Copernicus Land Management, 2018, 100m) and selected the class “sandy beaches” (code 62111) as the coastal area for our model predictions. Within this area, environmental predictor variables (annual sand dynamics and elevation relative to SHW) were extracted at 1 m resolution using R (*terra*). Predictor values were standardized using the same scaling parameters as applied during model fitting. Spatial predictions were based on environmental conditions for the year 2023, assuming these are representative of recent coastal conditions.

The ZINB abundance model yields an expected local abundance/cover value, *μ*. To translate expected abundance/cover into discrete simulated values, we sampled from the fitted zero-inflated negative binomial distribution using the predicted mean abundance/cover *μ* and the species-specific overdispersion parameter. These simulations therefore retained both the excess-zero structure and the overdispersed positive abundance values of the fitted model (equation 5).

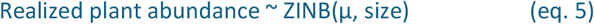

where *μ* is the predicted mean abundance and *size* is the species-specific overdispersion parameter estimated by the model. These simulations provide plausible realizations of potential realized plant abundance under the predicted environmental conditions.

### Sand accumulation

We quantified species-specific sand accumulation using independent field measurements of embryo dunes formed by the same four species (*Cakile maritima*, *Calamagrostis arenaria*, *Elymus farctus*, and *Salsola kali*). During three consecutive summers (2023, 2024 and 2025), we surveyed the dry beach for embryo dunes on the mid coast (Oostende) and west coast (Nieuwpoort-De Panne). We measured the volume sand captured by vegetation using the LiDAR sensor on an iPhone 13 Pro. Using the 3D Scanner App (SRL, 2024). We generated high-density point clouds of each embryo dune at centimeter resolution. Point clouds were exported as .las files, along with scaled orthophoto.

Point cloud processing was performed in CloudCompare (v2.13.2, GPL software (2026). Individual embryo dunes were extracted by cleaning and clipping each cloud, and multiple dune-clouds were separated into individual units. To avoid bias due to misalignment, cloud orientations were flattened using PCA, and coordinates were normalized by setting the origin. Sand volumes, surface areas, and maximum height were calculated using the *Volume 2.5D* tool on a 0.01m grid, with interpolation of small gaps where necessary. Vegetation cover associated with each embryo dune was quantified from orthophotos using Fiji (Schindelin et al., 2012). After setting the spatial scale, vegetation patches were manually delineated and their surface area calculated. A visualization of this workflow is available in S.1.4-S.1.6. To quantify the relationship between vegetation and sand accumulation, we fitted species-specific linear models relating sand volume (m³) to vegetation cover (m²). The observations for plant cover were root-transformed to meet the assumptions for a linear model. Linear models achieved highest R² relative to alternative exponential models. We modelled the amount of sand (m³) captured per m² (mean vertical accretion - m) for each species separately to quantify the efficiency by which vegetation cover promotes sand trapping (equation 6).

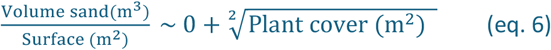

The regression was constrained to pass through the origin (intercept = 0), reflecting the assumption that vegetation is required for plant-induced sand accumulation, and that background sand deposition is not considered in this analysis.

Estimated slopes from these models were used to upscale sand accumulation efficiency to potential accumulated sand volumes along the coastline using the modelled potential realized abundance of each species. Average vegetation cover per species was calculated from field observations (orthophotos) and used to derive a mean sand accumulation per plant (equation 7).

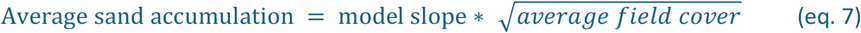

Total accumulation was then estimated by combining species-specific accumulation rates with spatial predictions of potential realized abundance. To account for uncertainty in spatial organization of vegetation, we considered two contrasting scenarios:

1. Individual-based scenario: plants function as isolated units, each contributing independently to sand accumulation based on the average observed cover per plant (equation 8).

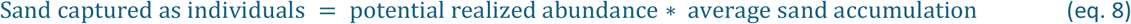

2. Patch-based scenario: plants form continuous patches, in which sand accumulation scales with the combined vegetation cover (equation 9).

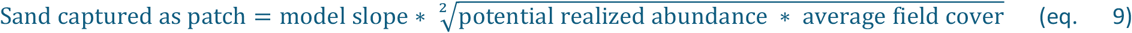

These calculations assume that relationships between vegetation cover and sand accumulation observed at local sites are representative across the Belgian coastline. Given limited sample sizes for some species (notably *C. arenaria* and *S. kali*), resulting estimates should therefore be interpreted as indicative of potential magnitudes rather than precise values. A comprehensive, visual overview of the statistical methodology and post-processing is illustrated in Fig. 2.

**Fig. 2.**
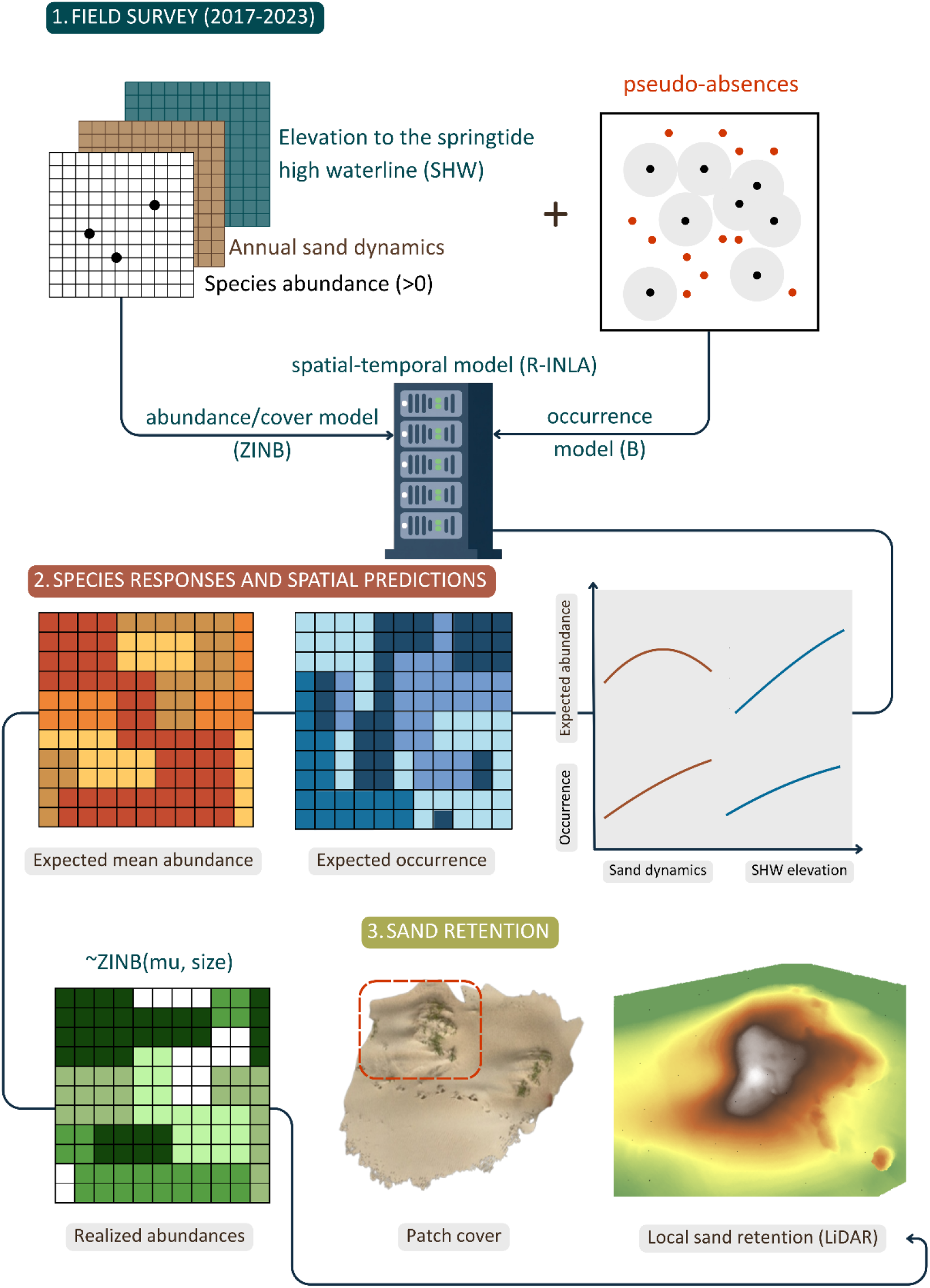
Overview of the data-processing and modelling workflow. Georeferenced field observations of pioneer strandline plants were converted into two complementary model responses: occurrence, based on presences and ecologically defined pseudo-absences, and abundance/cover, based on a rasterized response with positive values in occupied cells and zeros in unoccupied cells. Both responses were modelled using LiDAR-derived elevation relative to springtide high water (SHW) and annual sand dynamics within low-disturbance reference zones and then projected across the Belgian coastline. Predicted abundance/cover was subsequently combined with field-measured vegetation cover–sand accumulation relationships to estimate potential embryo-dune sand retention.

## Results

### Surveyed plant distributions and model domain

In total we collected 21,861 geolocated observations of plant abundance across the full survey period, including 3506 *C. arenaria*, 9803 *C. maritima*, 7499 *E. farctus* and 1053 *S. kali*. Of these, 14718 observations were in the predefined regions of interest (ROIs) and were used for INLA model fitting, supplemented with 14,549 pseudo-absences. Field values for fitting (ROI) data ranged between [-0.99, 10.9] for SHW elevation and [-2.49, 2.65] for annual sand dynamics. Field values for pseudo-absences ranged between [-2.53, 15.7] for SHW elevation and [-2.78, 2.73] for annual sand dynamics. The 95% intervals of field data for each variable can be found in S.1.8.

### Abiotic controls on plant occurrence and abundance

Occurrence and abundance/cover responses were broadly consistent across 10 model iterations. Predicted occurrence probabilities were generally weakly affected by annual sand dynamics for most species (Fig. 3). The main exception was *Salsola kali*, for which occurrence reached a maximum at moderate sand accretion, approximately 0.50 m. In contrast, expected abundance/cover showed clearer responses to annual sand dynamics. Most species exhibited increasing expected abundance/cover under stronger annual sand burial. *Cakile maritima* showed an optimum around 0.50 m of annual sand accretion, whereas the predicted optima for *Elymus farctus* and *Calamagrostis arenaria* occurred beyond the range commonly observed in the field. The abundance/cover response of *Salsola kali* was comparatively flatter than that of the other species.

**Fig. 3.**
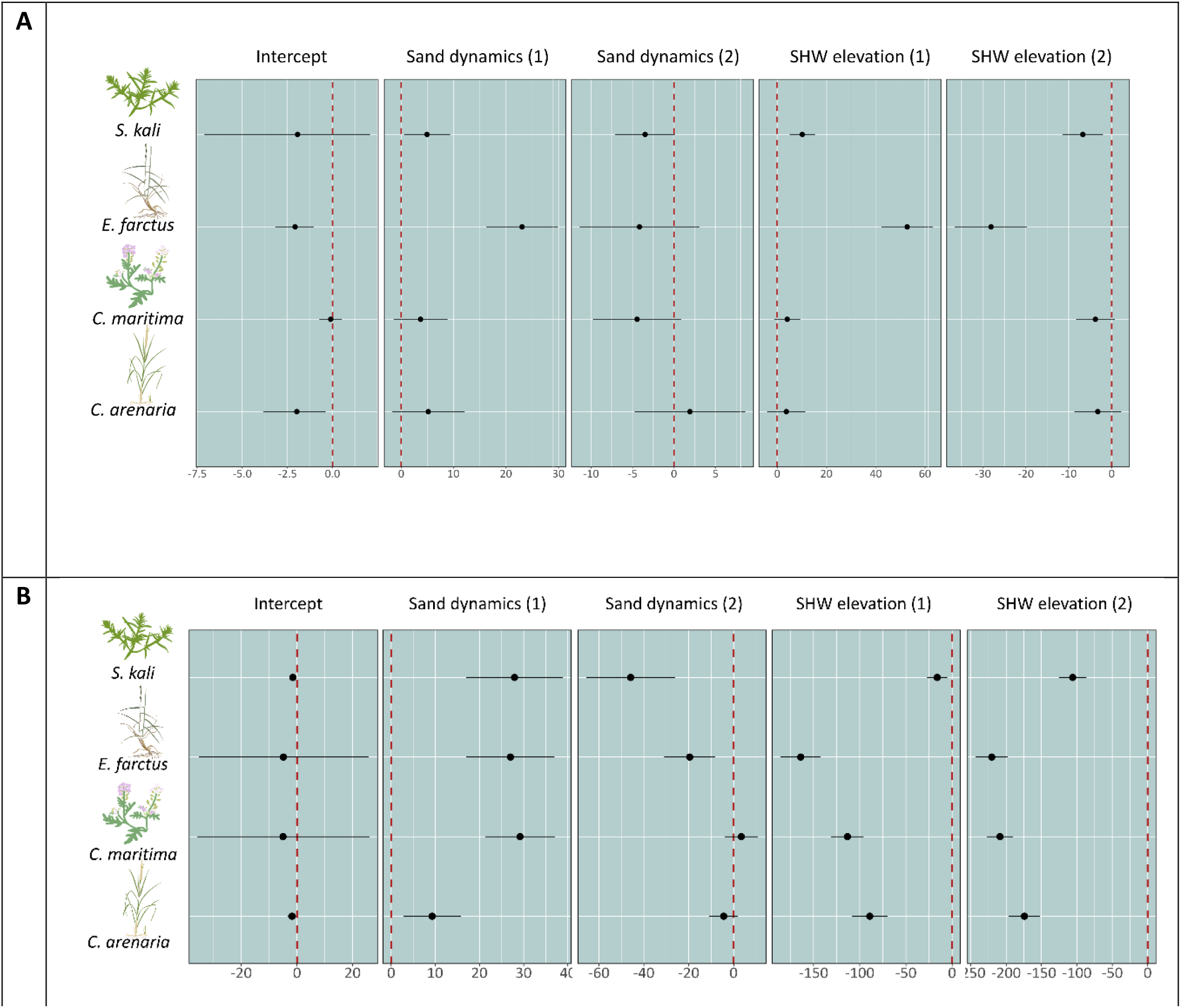
Summary of effect sizes across model iterations for (A) the abundance model and (B) the occurrence model. Posterior means, upper and lower bounds (95% CI) of the model parameters were averaged out for each species over all 10 model iterations. The black dots represent the mean of the posterior mean and visualize a distribution stretching to the left, indicating the mean of the lower CI bound, and to the right, indicating the mean of the upper CI bound. The red dotted line represents the zero-line; effect sizes are clearly determined when 0 is not included in the 95% credibility intervals of the posteriors

Posterior effect size estimates for the abundance/cover model were broadly consistent across model iterations (Fig. 3B). Annual sand dynamics showed a positive association with expected abundance/cover of *S. kali* and *E. farctus*, whereas effects for *C. maritima* and *C. arenaria* were weaker, with credibility intervals overlapping zero. The quadratic effect of sand dynamics was negative for S. kali, indicating a decline in expected abundance/cover at the highest levels of sand accretion, while quadratic effects for the other species were less clearly supported. Elevation relative to SHW was positively associated with expected abundance/cover for most species, with evidence for unimodal responses for *S. kali* and *E. farctus*. For *C. maritima* and *C. arenaria*, elevation effects were weaker and less clearly defined. Summary statistics of parameter estimates for both occurrence and abundance/cover models are provided in S.2.

The estimated spatial random fields, representing residual spatial autocorrelation not explained by the abiotic predictors, varied among species and years, but showed consistent patterns across model iterations (in S.2). Predicted relationships between occurrence probability, expected abundance/cover and the two environmental gradients were also consistent among iterations (Fig. 4). Across species, occurrence and abundance/cover generally increased with elevation relative to SHW. However, the abundance/cover response was least pronounced for *Calamagrostis arenaria* and *Salsola kali*. The fitted response curves suggested theoretical optima at high elevations relative to SHW, but these optima generally exceeded the range of values commonly observed in the field.

**Fig. 4.**
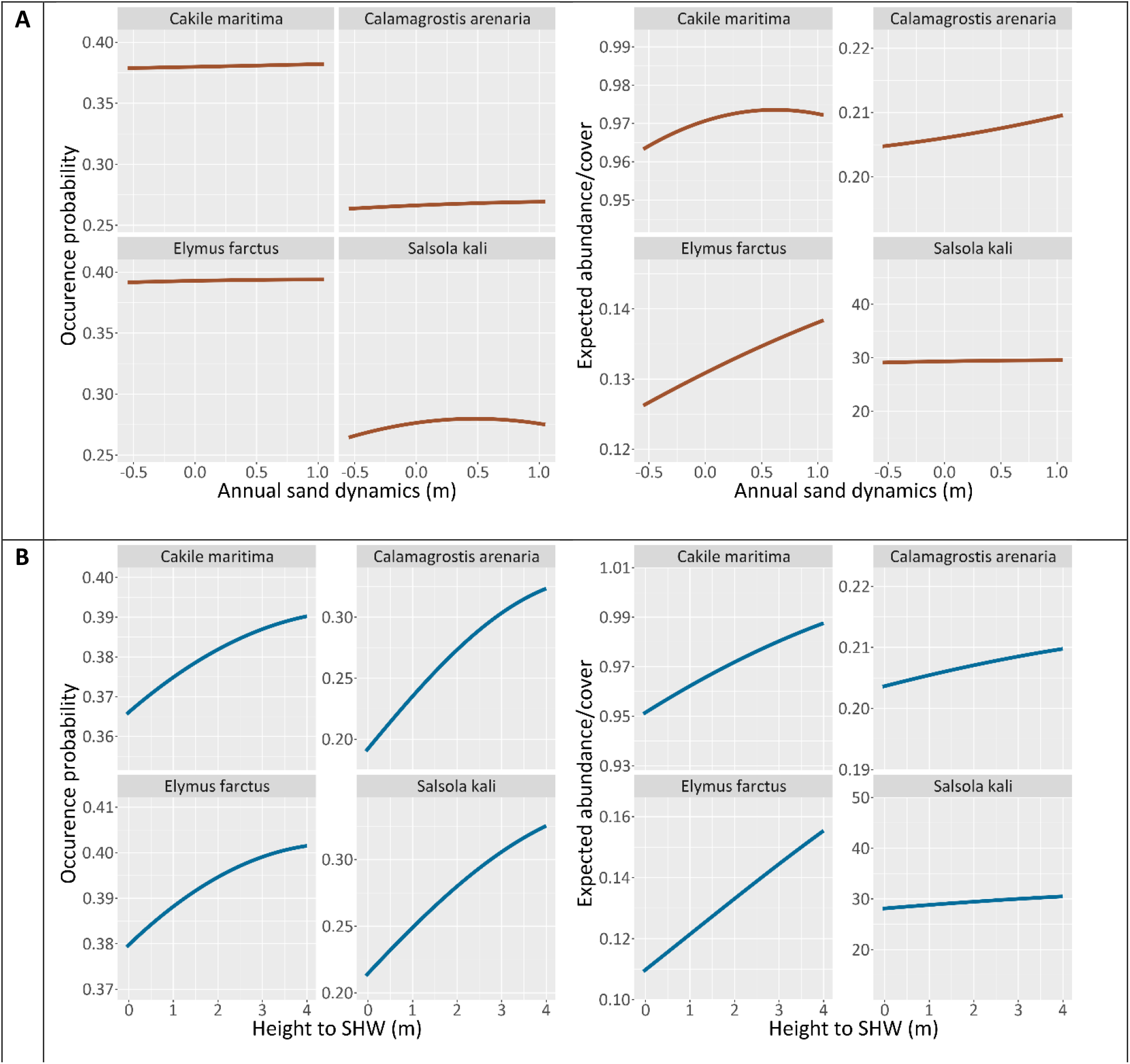
Predicted occurrence probability and expected abundance/cover across simulated values of annual sand dynamics (panel A) and elevation relative to SHW (panel B). Lines represent mean predictions across the 10 model iterations

### Coastal patterns of potential vegetation development

Spatial predictions revealed substantial heterogeneity in potential occurrence and expected abundance across the Belgian coastline. Since this heterogeneity was clear at both regional and local scales, we chose the municipality of De Panne as an illustrative example for the latter. This municipality features a wide, heavily used beach and is bordered by dune complexes to the west and east (De Westhoek and Zeepark-dunes; Fig. 5). Spatial occurrence and abundance/cover predictions for the entire coastline are presented in S.3.

**Fig. 5.**
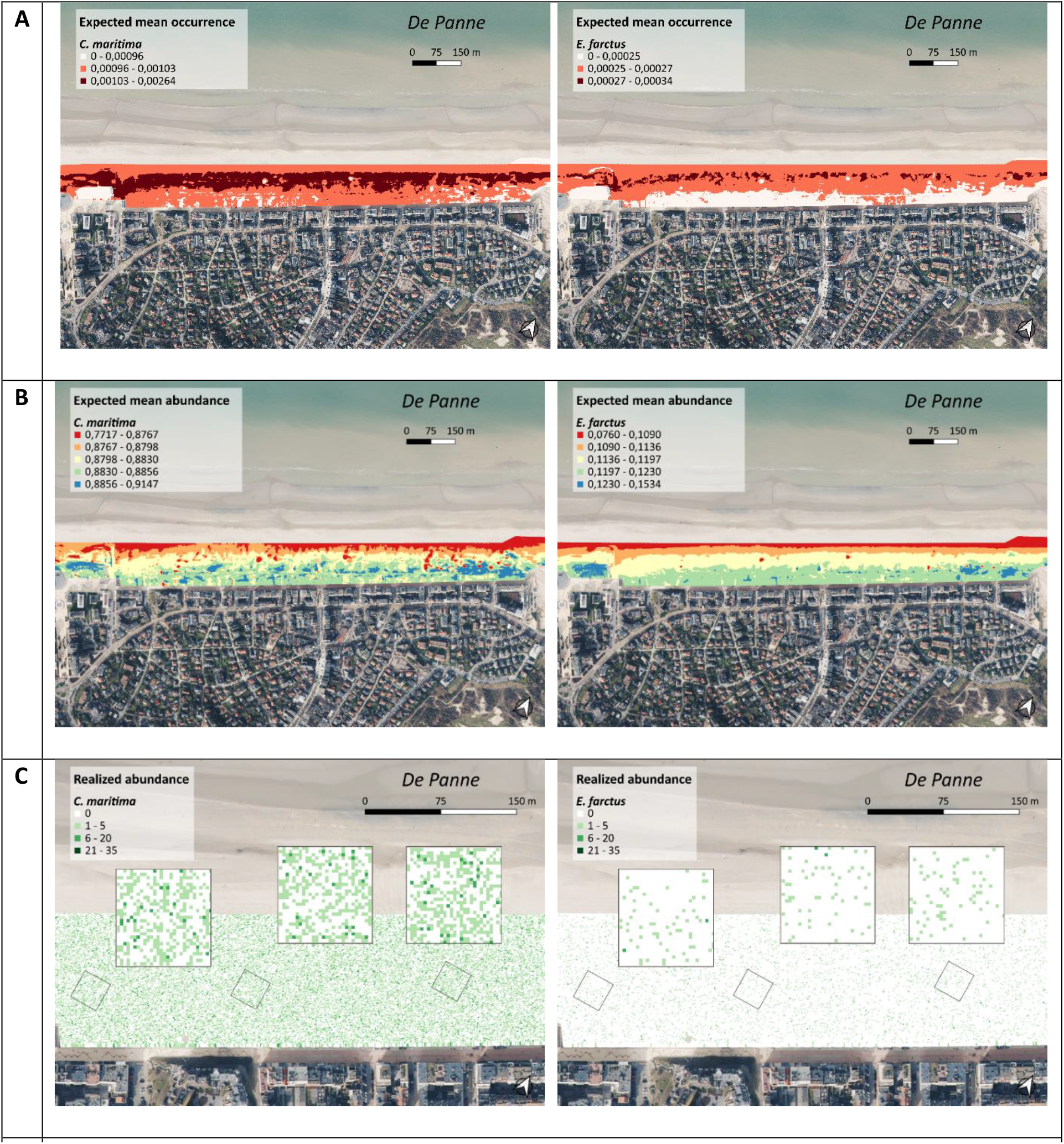
Spatial predictions for Cakile maritima and Elymus farctus on the beach of De Panne under low-disturbance reference conditions. Panel A: Predictions of the expected mean occurrences. Darker reds indicate higher occurrences. Panel B: Predicted mean abundances/cover. Red indicates low expected abundance/cover and blue indicates high expected abundance/cover. Panel C: Simulated abundance/cover for Cakile maritima and Elymus farctus in the De Panne area, illustrating stochastic variation around expected abundance/cover predictions.

First, predicted occurrences on the beach increased towards the sea for *Cakile*, *Elymus* and *Calamagrostis* (Fig. 5A). For *Salsola* however, occurrences did not exhibit a clear spatial pattern, as high and low occurrences were present along the entire beach strip.

Furthermore, expected abundance/cover was relatively high across the entire beach profile for *Cakile maritima*, indicating that abiotic conditions are broadly suitable for vegetation development by this species (Fig. 5B). However, local abundances tended to slightly increase towards the upper beach near the dike. *Elymus farctus* displayed a more pronounced cross-shore zonation, with mean abundances/cover increasing towards the upper beach in front of the dike. Predicted patterns for *Calamagrostis arenaria* were broadly comparable to those of *Elymus fractus*, with higher expected abundance/cover closer to the dike. In contrast, *Salsola kali* showed relatively high and spatially consistent expected abundance/cover across much of the beach compared with the other species.

At the coast-wide scale (in S.3), we detected areas with high predicted abundances both on urbanized beaches in front of dikes and on less disturbed beaches adjacent to dune systems. Under conditions comparable to the low-disturbance reference zones used for model calibration, expected mean abundance/cover also varied strongly among species and across the beach profile. These spatial patterns reflect species-specific responses to the abiotic gradients included in the model and should therefore be interpreted as potential abundance/cover under low-disturbance conditions. The contrast between these predictions and the low observed abundance of strandline vegetation on many urbanized beaches highlights the likely importance of non-abiotic constraints, such as trampling, beach cleaning and other forms of human disturbance.

Finally, to illustrate stochastic variability around the expected abundance/cover predictions, we simulated realized abundance values by sampling from the fitted abundance distribution using the predicted mean abundance/cover (μ). These simulations produced spatial patterns consistent with the mean predictions, while illustrating additional local variability in abundance/cover. *Cakile maritima* generally exhibited higher and more continuous abundance values, whereas *Salsola kali, Elymus farctus* and *Calamagrostis arenaria* showed lower and more patchy distributions (Fig. 5C). Full distributions of simulated abundance values are provided in S.3.

### Sand accumulation potential by embryo-dune vegetation

Species-specific relationships between vegetation cover and sand accumulation are shown in Fig. 6, with full model outputs provided in S.4. Across all species, vertical sand accumulation (expressed as sand volume per unit surface area) increased with vegetation cover, with the square root of plant cover showing a positive association with sand accumulation. Estimated regression slopes were broadly similar among *Cakile maritima* (0.159, SE=0.006, p<0.001) and *Elymus farctus* (0.155, SE=0.006, p<0.001), and slightly lower values for *Calamagrostis arenaria* (0.142, SE=0.010, p<0.001) and *Salsola kali* (0.121, SE=0.014, p<0.001), indicating comparable efficiencies of vegetation in promoting vertical sand accumulation.

**Fig. 6.**
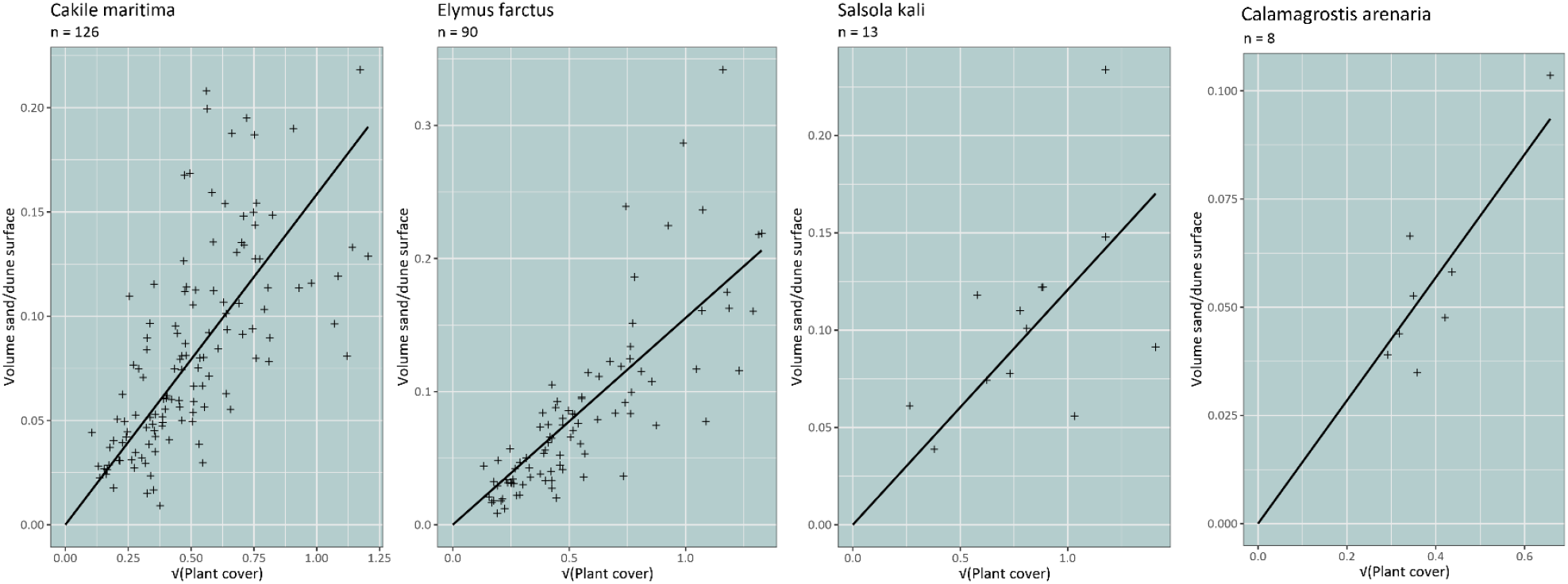
Relationship between mean vertical sand accumulation (m; volume per surface area) and vegetation cover (m²) for each species. Vegetation cover was square root transformed, and regression lines were fitted without intercept (intercept = 0).

Total sand accumulation was estimated by combining these species-specific relationships with spatial predictions of simulated abundance/cover. Predicted abundance values were converted to vegetation cover using species-specific average field cover estimates, after which sand accumulation was calculated for each species. Resulting sand volumes differed among species, with *Cakile maritima* contributing the largest share, followed by *Salsola kali*, *Elymus farctus*, and *Calamagrostis arenaria* (Table 1). Estimates depended on assumptions about the spatial organization of vegetation. Scenarios assuming aggregated vegetation into continuous patches produced higher total sand volumes than scenarios in which plants were treated as spatially dispersed units, reflecting the nonlinear scaling between vegetation cover and sand accumulation. These estimates assume that the locally measured relationships between vegetation cover and sand accumulation are representative across the coastline and should therefore be interpreted as indicative potential magnitudes rather than precise predictions.

**Table 1.** Predicted sand volumes for each species, as a patch or as distinct individuals, for the predicted number of plants on the Belgian coastline.

| Species | Sand volume (m<sup>3</sup>) |  |
| --- | --- | --- |
|  | Patch | Individuals |
| *C. arenaria* | 26033 | 29147 |
| *C. maritima* | 238833 | 386682 |
| *E. farctus* | 42090 | 56545 |
| *S. kali* | 69419 | 80095 |
| **Total volume (m<sup>3</sup>)** | **376375** | **552469** |

## Discussion

In this study, we combined niche modelling based on low-disturbance occurrences with field-based remote sensing to assess the potential establishment and sand accumulation capacity of four pioneer dune plant species along the highly urbanized Belgian coast. Our results indicate that abiotic conditions along large parts of the coastline are suitable for plant establishment. The contrast between predicted potential abundance/cover and the generally low observed vegetation cover on many urbanized beaches suggests that factors not included in the abiotic models, particularly human disturbance, strongly constrain the realized distribution of strandline vegetation. In addition, our analyses show that, once established, pioneer vegetation can contribute substantially to sand accumulation through embryo dune formation. The total volumes of sand potentially retained by these bio-geomorphic processes are of a similar order of magnitude to the sediment volumes annually supplied to the system to sustain dune development. Rather than replacing engineered interventions, these results highlight the potential role of strandline vegetation in enhancing the efficiency with which nourished sediment is retained within the coastal system.

### Boundary conditions for pioneer beach plants

Our models indicated that large parts of the Belgian coastline provide suitable abiotic conditions for the settlement and persistence of pioneer strandline species (*Cakile maritima*, *Elymus farctus*, *Calamagrostis arenaria* and *Salsola kali*). In low-disturbance conditions (absence of selective beach cleaning and excessive tourist trampling), conditions controlled by sediment dynamics and elevation generally permit plant establishment. This suggests that, across much of the Belgian coast, the abiotic conditions represented by elevation relative to SHW and annual sand dynamics are not the primary constraints on potential occurrence of strandline vegetation under low-disturbance conditions.

While all species showed broadly similar responses to abiotic gradients, interactions among strandline species are likely weak or highly context-dependent rather than absent. Recent work demonstrates that dune-building grasses can simultaneously facilitate and compete depending on spatial scale, with pioneer species modifying environmental conditions at small scales while competing for space at larger scales (Lammers et al., 2024). Under the highly dynamic and stressful conditions of the beach, abiotic filtering likely dominates, but biotic interactions may still contribute to local spatial structure.

A key result of our study is that annual sand dynamics affect occurrence and abundance/cover differently. Occurrence probabilities were relatively insensitive to annual sand dynamics for most species, suggesting that sand accretion is not the main determinant of whether plants are present at the end of the growing season. In contrast, expected abundance/cover generally increased with sand accretion, indicating that sediment dynamics more strongly influence post-establishment growth and patch development than initial establishment itself. At first sight, this appears to contrast with experimental evidence showing that burial can constrain seed germination (Bonte et al., 2021; Lee, 1985). This apparent discrepancy can be explained by distinguishing between life stages. Burial may limit germination or early seedling survival, but once plants have established, moderate sand accretion can stimulate vertical growth, cover expansion and clonal spread (Davy et al., 2006; Maun, 2009; van Egmond et al., 2019). Because our surveys were conducted late in the growing season and abundance was derived from cover-based classes, the abundance/cover model primarily reflects post-establishment vegetation development rather than recruitment success. This interpretation is particularly relevant for clonal dune-building grasses such as *Calamagrostis arenaria*, for which moderate burial can stimulate vertical growth and clonal expansion once seedlings have established (Bonte et al., 2021; Lammers et al., 2023; Reijers et al., 2021).

The degree to which beach plants are subject to seawater inundation and salt spray (Davy et al., 2006) is defined by their relative position to the springtide water level. For all species, predicted abundance and potential occurrence increased with higher elevations above the springtide level, indicating a trade-off between moisture availability closer to the waterline and increasing physiological stress due to salt exposure (Debez et al., 2018). Species-specific optima were mainly theoretical and exceeding their natural range of field observations. Differences in the strength of these responses likely reflect variation in physiological tolerance to salinity and drought, as well as differences in life-history strategies.

### Spatial suitability and distribution

Predicted occurrence and expected abundance/cover revealed clear species-specific spatial patterns, reflecting differences in establishment capacity, growth form and responses to cross-shore abiotic gradients. *Cakile maritima* for instance showed a coast-wide high suitability for plant establishment in both urban and low-disturbance areas, as predictions illustrated large areas of high expected abundance/cover across the entire east-west axis. This broad tolerance is consistent with its life history as a short-lived strandline annual, capable of rapid recruitment and vegetation development when disturbance is low and abiotic conditions are favorable (Michael, 1970a, 1970b). Notably, dense populations of *Cakile* were observed during the COVID-19 lockdown period (S.1.1), when human disturbance was minimal, supporting the interpretation that suitable abiotic conditions alone permit widespread establishment when disturbance pressure is reduced.

Furthermore, *Calamagrostis arenaria* and *Elymus farctus* showed generally low predicted abundances/cover across the beaches in heterogeneous mosaic-like spatial patterns covering different types of beaches, however with higher values restricted to specific locations on the west and east coast for *Calamagrostis* and *Elymus,* respectively. This pattern probably reflects a restricted establishment window driven by constraints on seed or rhizome arrival and germination along the beach profile. Both species also exhibited the highest occurrence chances on the eastern part of the coast, despite in much higher magnitudes for marram grass. Still, both grasses had similar predicted spatial distributions of individuals on the beach showing low, patchily distributed expected abundance/cover. This is remarkable since *Calamagrostis* and *Elymus* have different growth forms and thus spatial patterns as dune-building species. Likely, since marram grass is a dominant foredune species (Gray, 1985), the species might encounter suboptimal conditions at the exposed strandline zone on the dry beach, resulting in more spatially dispersed organization similar to *Elymus*, rather than its high-dense, patchy distribution typical in the foredunes (Lammers et al., 2023).

*Salsola kali* illustrates the added value of distinguishing occurrence from abundance/cover. Although predicted occurrence was high across much of the coastline, expected abundance/cover was higher on the west coast than elsewhere. This indicates that abiotic conditions may allow widespread establishment, while local vegetation development remains spatially restricted. This pattern is consistent with field observations of *Salsola* as a non-clonal annual that can disperse seeds over relatively long distances by tumbleweed movement (Ayres et al., 2008) but does not necessarily form dense local cover everywhere it occurs.

### Sand accumulation through embryonic dunes

Our results confirm a positive relationship between vegetation cover and local sand accumulation, indicating that pioneer vegetation promotes embryo-dune formation through its effects on aeolian sediment trapping. Across all species, sand accumulation (i.e., sand volume per unit surface area) increased with cover. Differences in sand accumulation efficiency among species were relatively small, with slightly higher values for *Cakile maritima* and *Elymus farctus* compared to *Calamagrostis arenaria* and *Salsola kali*. These differences likely reflect variation in plant architecture and growth dynamics, although they remain modest relative to the strong effect of vegetation cover itself.

The comparatively lower sand accumulation efficiency of marram grass likely reflects its poor performance, and subsequent low cover, on the higher beach. While marram grass is known to capture large volumes of sand in foredune environments through rapid clonal expansion under sustained burial (Bonte et al., 2021; Seabloom et al., 2013; Zarnetske et al., 2015), individuals on the beach are primarily seedlings exposed to elevated levels of salt stress and sediment disturbance. As a result, their contribution to sand accumulation remains limited compared to their well-known engineering capacity in established dune systems.

When scaled to the coastline under low-disturbance assumptions, our results suggest that pioneer beach vegetation could retain substantial volumes of sand through embryo-dune formation. These volumes are of a similar order of magnitude to the sediment inputs from nourishment required to sustain ongoing dune development, emphasizing the potential importance of vegetation-mediated processes in retaining sand within the coastal system. Rather than replacing engineered interventions that provide the needed upper beach conditions, these findings highlight that embryo dunes may enhance the efficiency with which externally supplied sediment is captured and retained along the beach.

### Implications for management

The distinction between occurrence and abundance/cover has practical management relevance. If abiotic conditions already allow occurrence across large parts of the coast, then management interventions do not necessarily need to create entirely new physical habitat everywhere. Instead, reducing disturbance may allow existing suitable areas to transition from potential occurrence to substantial vegetation cover, thereby increasing embryo-dune formation and sediment retention. Simple management interventions such as limited trampling and manual beach cleaning may therefore be sufficient to promote vegetation establishment and enhance strandline biodiversity. The designation of specific beach zones, like in some larger cities to create rest zones for seals, provides a clear example where reduced disturbance has coincided with rapid development of strandline vegetation. These effects can emerge within a single growing season, highlighting the potential for low-cost and rapidly effective nature-based interventions.

Promoting the natural establishment of pioneer dune species and the development of embryo dunes may contribute directly to economic benefits through the retention of sand within the coastal system. This sand is then not eroded back to the sea or inducing nuisance on dikes and promenades (Van der Biest et al., 2017). Rather than substituting for engineered interventions, these vegetation-driven processes may act as a complementary mechanism that supports coastal protection strategies (Van der Biest et al., 2017).

Our modelling approach should be interpreted as an assessment of the first step in dune development: where pioneer vegetation can establish and how much sand may initially be trapped by this early vegetation. It does not predict how dunes will subsequently grow, merge, erode or develop into more permanent foredune structures. This distinction is important because the establishment of vegetation is not only a response to existing beach conditions but can also modify those conditions. Once plants trap sand and embryo dunes start to form, local elevation increases, exposure to marine inundation may decrease, and sediment transport across the upper beach may be redirected. As a result, the beach surface on which future vegetation establishes may differ from the abiotic template used in our models.

Embryo dunes therefore represent a transitional stage between the open beach and the foredune system. If pioneer vegetation persists, the small dunes it creates may provide more favorable conditions for later dune-building species, including *Calamagrostis arenaria*. In the longer term, marram grass may expand clonally from these incipient dunes, trap additional sand, and promote the development of larger and more stable foredune structures (Bonte et al., 2021). Our estimates of sand accumulation should therefore be interpreted as the potential contribution of early strandline vegetation to the initiation of this process, rather than as predictions of long-term dune growth or coastal morphology.

In coastal settings characterized by a positive sediment balance, such vegetation-driven accumulation may contribute to increasing coastal safety by promoting dune growth and stability. In contrast, under more sediment-limited or erosive conditions, increased sand capture by vegetation may lead to a redistribution rather than a net gain of sediment, potentially creating a dynamic equilibrium between beach and dune systems. In such cases, vegetation may still play a key role by reducing sediment losses through alongshore transport and retaining sand within the coastal compartment, even if net accretion is limited. These processes may also alter sediment availability on the beach itself, as increased capture of aeolian sand may reduce the supply of sediment available for redistribution across the beach profile (Davidson-Arnott et al., 2018; Delgado-Fernandez & Davidson-Arnott, 2011). Such feedbacks could have implications for beach morphology and sediment budgets, potentially affecting coastal dynamics beyond the initial stages of dune formation. Understanding these longer-term interactions between vegetation, sediment transport, and coastal morphology therefore requires further multidisciplinary investigation.

## Acknowledgements

This work was supported by a predoctoral fellowship from the Research Foundation - Flanders (FWO) awarded to Charlotte Taelman (grant number 1SH0I24N). Dries Bonte received support by a UGent BOF research grant BOF/24J/2021/066 and the European Union’s Horizon Europe research and innovation program under grant agreement No739 101135410 – The DuneFront Project. The computational resources (Stevin Supercomputer Infrastructure) and services used in this work were provided by the VSC (Flemish Supercomputer Centre), funded by Ghent University, FWO and the Flemish Government – department EWI. Guidance in the R-INLA coding was provided by the helpful community of the R-INLA forum.

## Author contributions

Charlotte Taelman and Sam Provoost conceived the ideas and designed methodology. Charlotte Taelman and Sam Provoost collected the data in the field. Charlotte Taelman analyzed the data. Charlotte Taelman led the writing of the manuscript. All authors contributed critically to the drafts and gave final approval for publication.

## Data availability

The raw data and coding scripts will be made available on Zenodo upon publication (10.5281/zenodo.21357573).

