## Supplementary materials for "From strandline vegetation to embryo dunes: natural sand retention along a heavily urbanized coast"

### S.1 Fieldwork and methodology

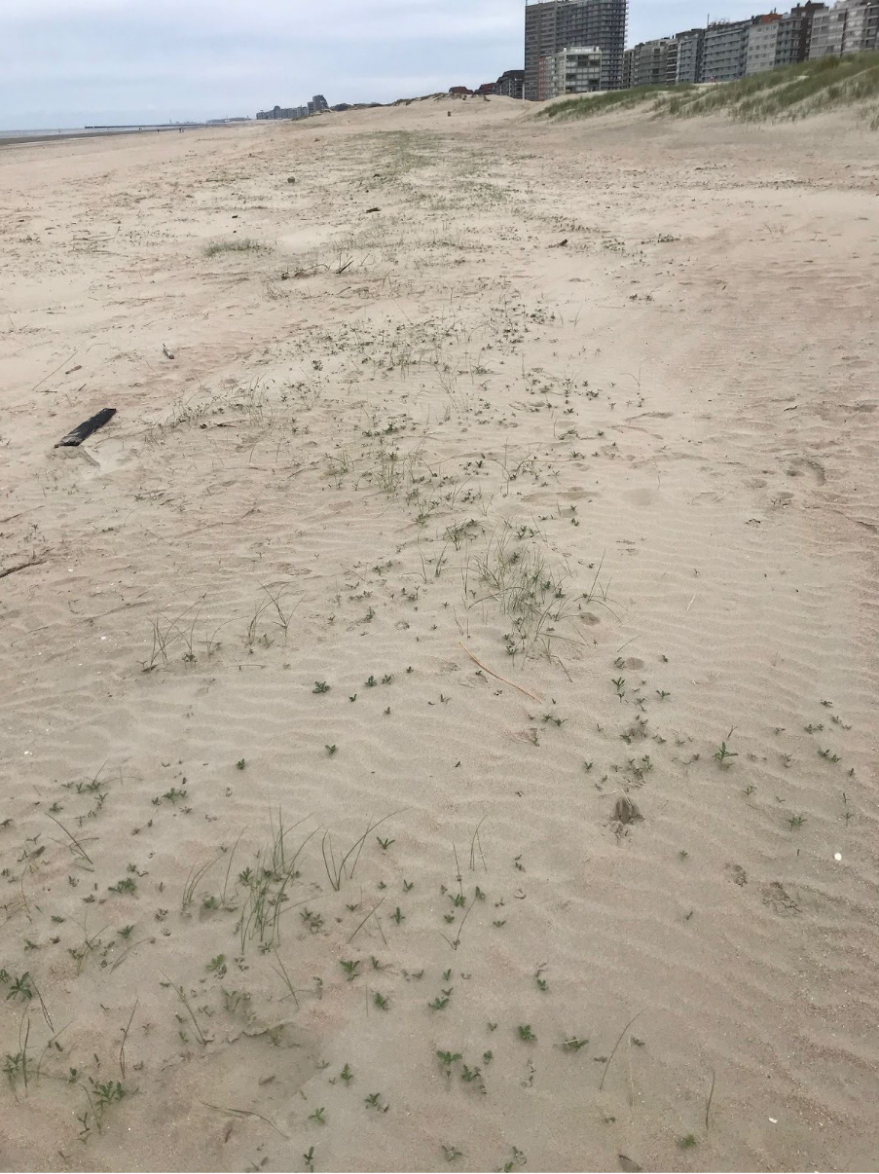

Fig. S.1.1. Beach during Covid19 pandemic. The beach of Oostduinkerke (03/05/2020) full of Cakile seedlings during the Covid19-pandemic. Photo taken by Ann-Katrien Lescrauwaet.

Table S.1.2. Conversion table for area and abundances of plants. In the field, observed abundances of beach plants were categorized into classes with a specific code, referring to a median abundance value. To properly assign the correct class to an observed plant patch, we made a visual estimation of the radius (m) and surface (m²) of the patch. This method was used for all four study species.

| Code | Amount of individuals | Median | Surface (m²) | Radius of circle (m) |
| --- | --- | --- | --- | --- |
| a | 1 | 1 | < 1 | 1 |
| b | 2-5 | 3 | 1-5 | 1 |
| c | 6-25 | 12 | 5-25 | 2 |
| d | 26-50 | 35 | 25-50 | 3 |
| e | 51-500 | 200 | 50-500 | 8 |
| f | 501-5000 | 2000 | 500-5000 | 25 |
| g | >5000 | 7500 | >5000 | 49 |

Table S.1.3. Overview of LiDAR surveys.

| Year of sampling | Date | Resolution |
| --- | --- | --- |
| 2017 | 26/05/2017 | 1m |
|  | 06/11/2017 | 1m |
| 2018 | 17/04/2018 | 1m |
|  | 06/11/2018 | 1m |
| 2019 | 20/04/2019 | 1m |
|  | 29/10/2019 | 1m |
| 2020 | 10/04/2020 | 1m |
|  | 18/11/2020 | 1m |
| 2021 | 28/04/2021 | 1m |
|  | 23/02/2022 | 1m |
| 2022 | 17/04/2022 | 1m |
|  | 08/02/2023 | 1m |
| 2023 | 17/06/2023 | 1m |
|  | 16/01/2024 | 0.25m |

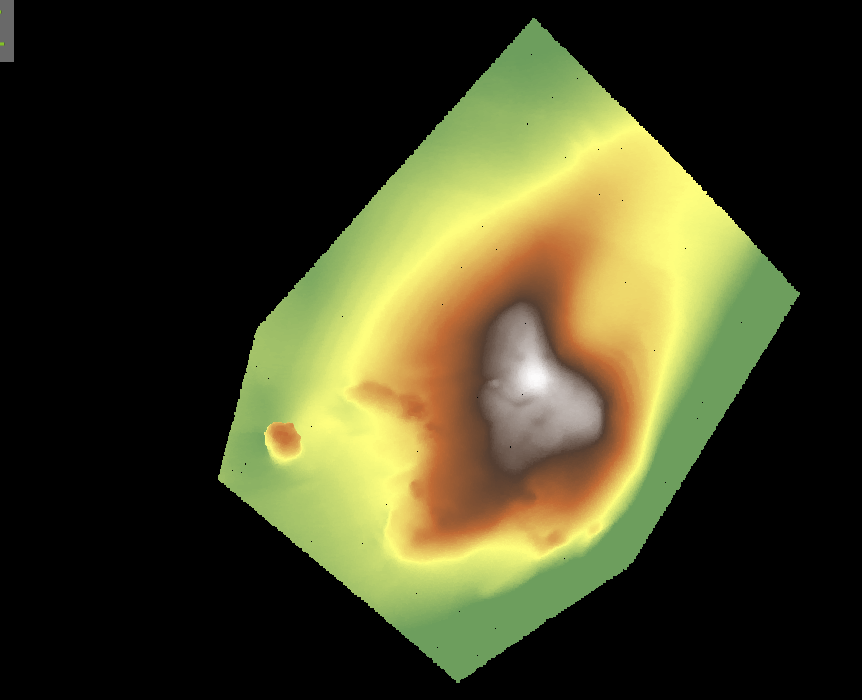

Fig. S.1.4. iPhone LiDAR scan in CloudCompare. The LiDAR scan of a Cakile-dune in the field. The terrain height is indicated by colors (green is low, grey is high). The scan was cropped to isolate the dune including the vegetation and reduce noise from the surrounding region. Volumes and surfaces were calculated in CloudCompare.

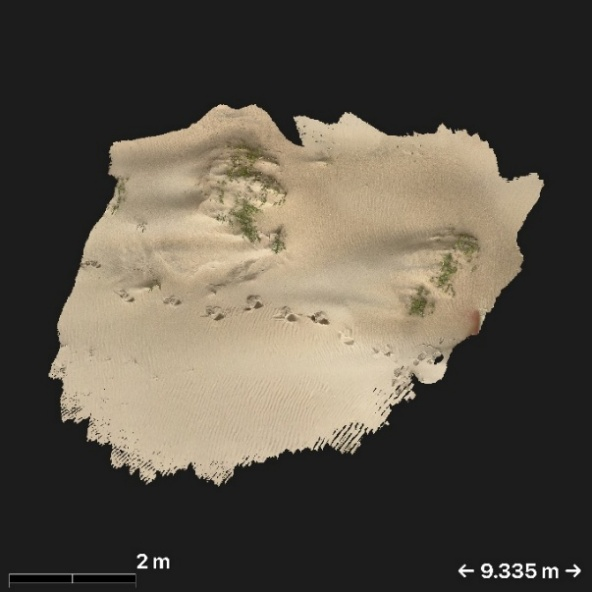

Fig. S.1.5. A scaled orthophotomosaic of the LiDAR scan.

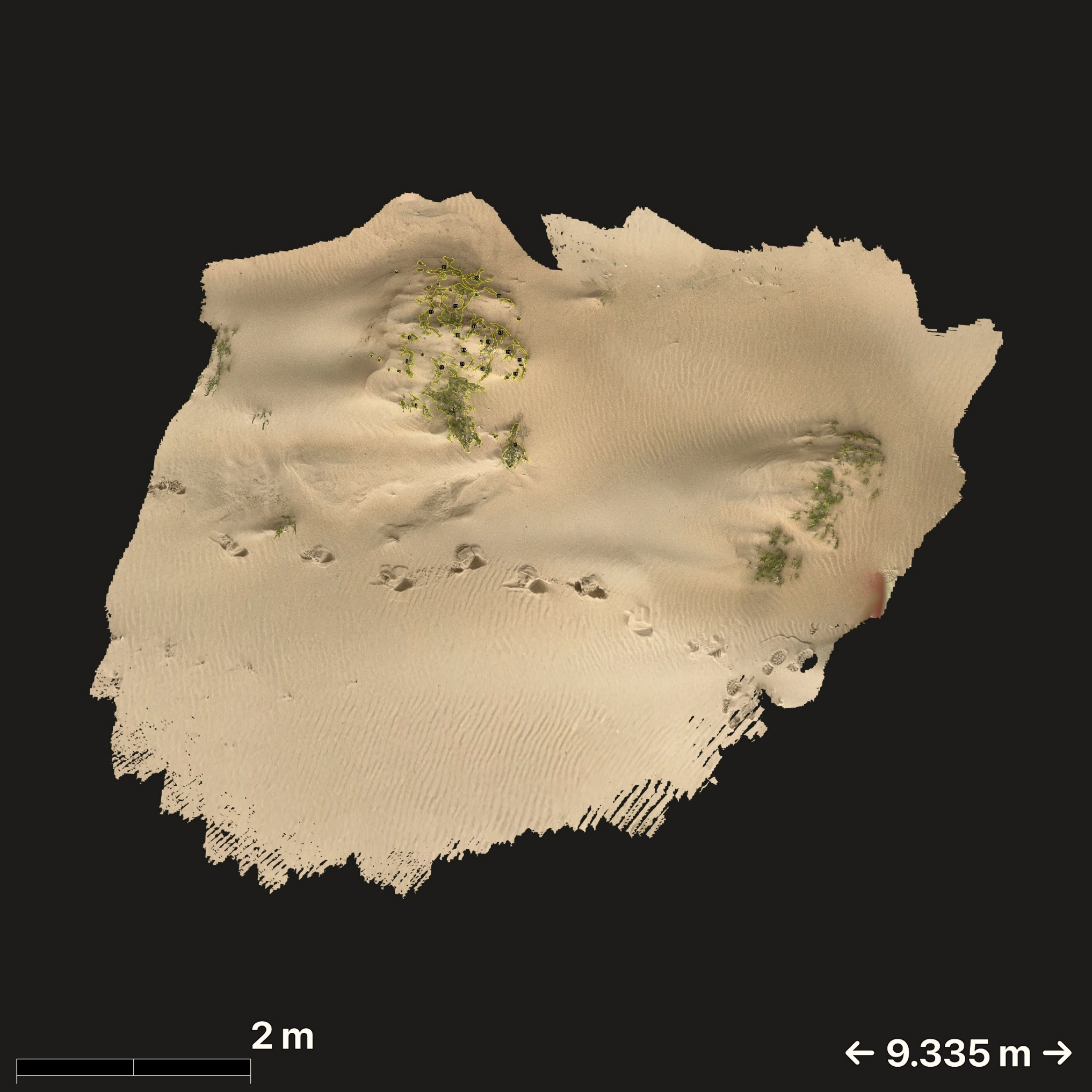

Fig. S.1.6. Yellow polygons indicating the edges of the vegetation in Fiji. We manually drew polygons around the edges of the vegetation as accurately as possible. These polygons were scaled (using the scale on the orthophoto), and their inside surfaces were summed.

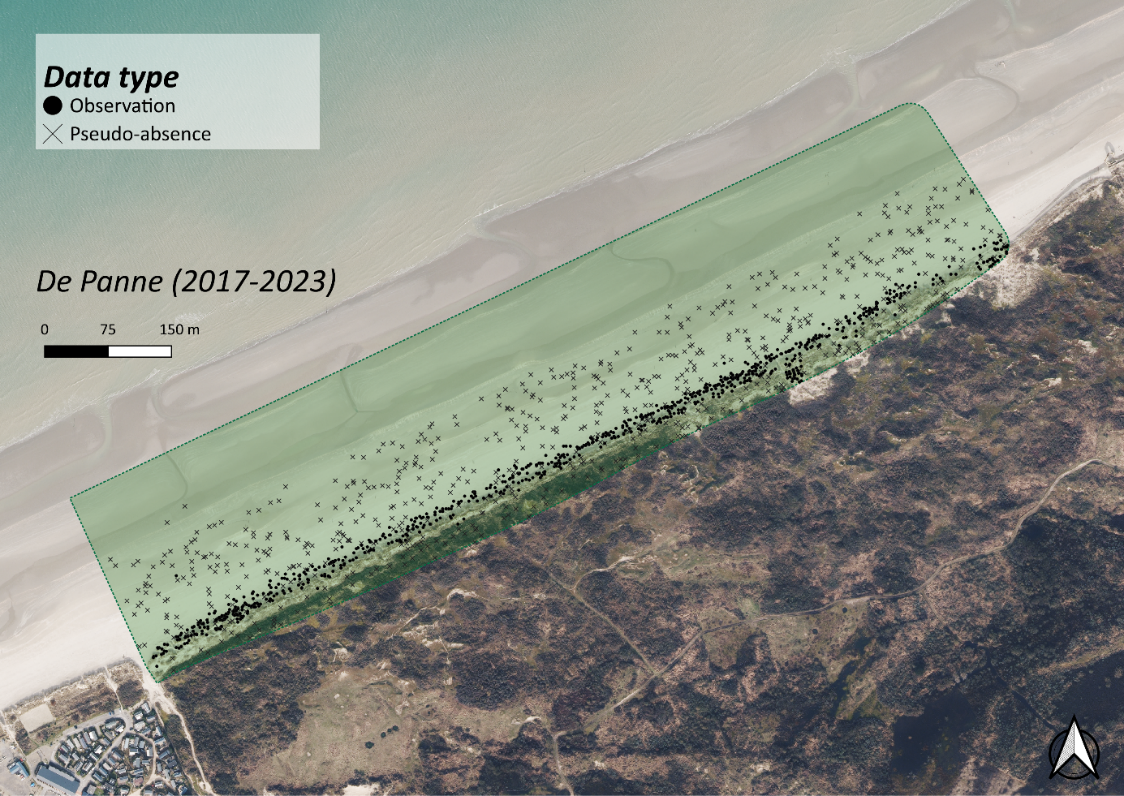

Fig. S.1.7. Map indicating observed data and generated pseudo-absence data over the entire sampling period.

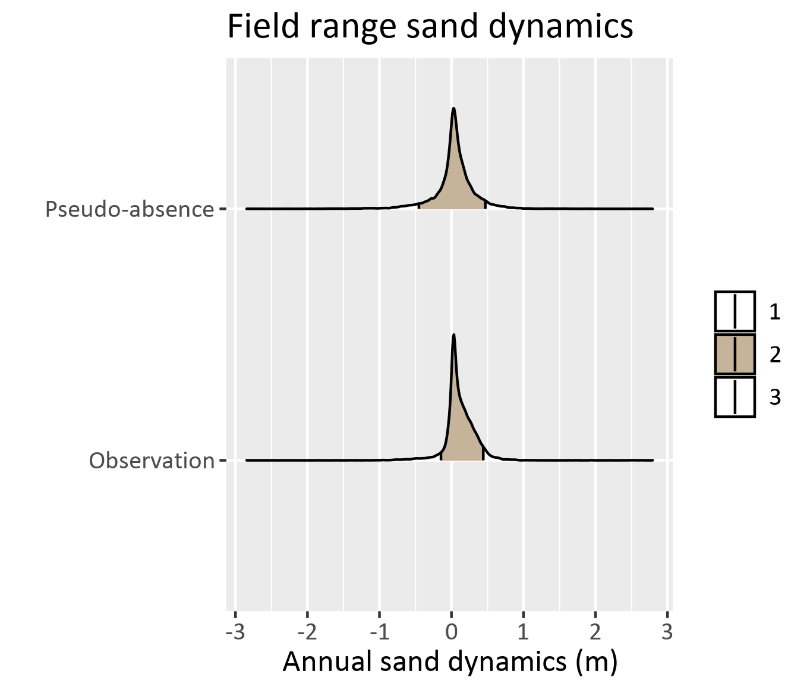

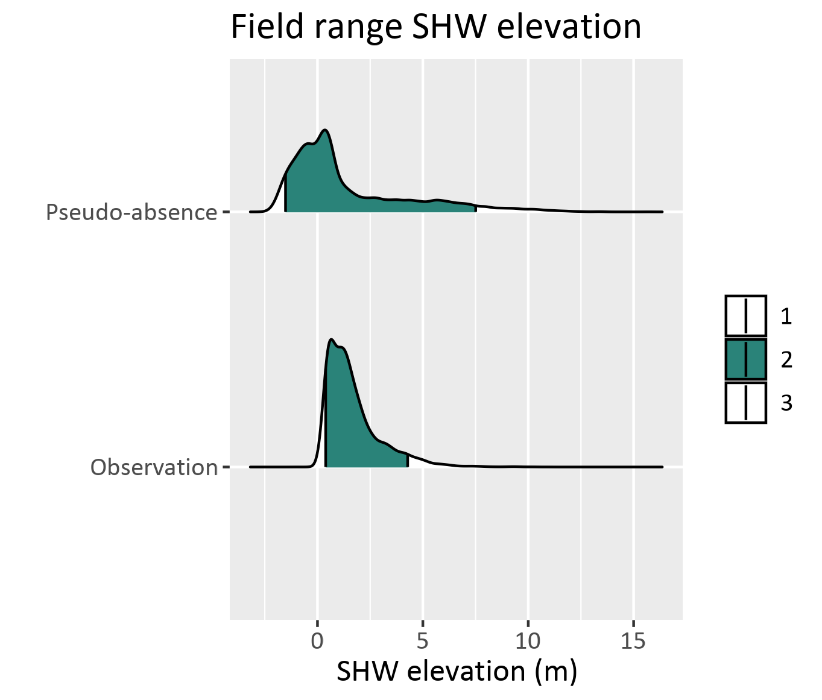

Fig. S.1.8. Field ranges of annual sand dynamics (left) and elevation to the SHW (right), together with the 90% intervals.

### S.2 INLA model

Table S.2.1. Means of the posterior effect sizes of the abundance/cover model.

| **Species** | **Variable** | **Mean(mean)** | **Mean(low)** | **Mean(up)** |
| --- | --- | --- | --- | --- |
| Calamagrostis arenaria | Intercept | -1.98 | -3.84 | -0.38 |
|  | Sand dynamics (1) | 5.16 | -1.71 | 12.12 |
|  | Sand dynamics (2) | 1.91 | -4.72 | 8.60 |
|  | Springtide elevation (1) | 3.71 | -4.08 | 11.44 |
|  | Springtide elevation (2) | -3.25 | -8.70 | 2.21 |
| Cakile maritima | Intercept | -0.10 | -0.73 | 0.52 |
|  | Sand dynamics (1) | 3.69 | -1.43 | 8.83 |
|  | Sand dynamics (2) | -4.44 | -9.76 | 0.88 |
|  | Springtide elevation (1) | 4.09 | -1.13 | 9.32 |
|  | Springtide elevation (2) | -3.80 | -8.31 | 0.72 |
| Elymus farctus | Intercept | -2.07 | -3.17 | -1.03 |
|  | Sand dynamics (1) | 23.04 | 16.24 | 29.89 |
|  | Sand dynamics (2) | -4.16 | -11.37 | 3.05 |
|  | Springtide elevation (1) | 52.73 | 42.36 | 63.16 |
|  | Springtide elevation (2) | -28.12 | -36.57 | -19.73 |
| Salsola kali | Intercept | -1.94 | -7.11 | 2.08 |
|  | Sand dynamics (1) | 4.92 | 0.57 | 9.33 |
|  | Sand dynamics (2) | -3.48 | -7.09 | 0.12 |
|  | Springtide elevation (1) | 10.18 | 5.06 | 15.40 |
|  | Springtide elevation (2) | -6.72 | -11.48 | -2.01 |

Table S.2.2. Means of the posterior effect sizes of the occurrence model.

| **Species** | **Variable** | **Mean(mean)** | **Mean(low)** | **Mean(up)** |
| --- | --- | --- | --- | --- |
| Calamagrostis arenaria | Intercept | -1.77 | -3.46 | -0.15 |
|  | Sand dynamics (1) | 9.28 | 2.80 | 15.75 |
|  | Sand dynamics (2) | -4.42 | -10.83 | 1.98 |
|  | Springtide elevation (1) | -89.12 | -108.19 | -70.08 |
|  | Springtide elevation (2) | -174.18 | -196.17 | -152.22 |
| Cakile maritima | Intercept | -5.03 | -35.77 | 25.96 |
|  | Sand dynamics (1) | 29.18 | 21.34 | 37.01 |
|  | Sand dynamics (2) | 3.42 | -4.04 | 10.90 |
|  | Springtide elevation (1) | -113.28 | -131.20 | -95.79 |
|  | Springtide elevation (2) | -208.66 | -227.34 | -189.96 |
| Elymus farctus | Intercept | -4.88 | -35.30 | 25.78 |
|  | Sand dynamics (1) | 27.00 | 17.02 | 36.98 |
|  | Sand dynamics (2) | -19.56 | -30.95 | -8.18 |
|  | Springtide elevation (1) | -163.88 | -186.02 | -142.42 |
|  | Springtide elevation (2) | -220.36 | -242.92 | -197.75 |
| Salsola kali | Intercept | -1.49 | -2.68 | -0.47 |
|  | Sand dynamics (1) | 27.92 | 16.95 | 38.92 |
|  | Sand dynamics (2) | -45.88 | -65.69 | -26.13 |
|  | Springtide elevation (1) | -16.13 | -27.04 | -5.23 |
|  | Springtide elevation (2) | -105.96 | -125.57 | -86.46 |

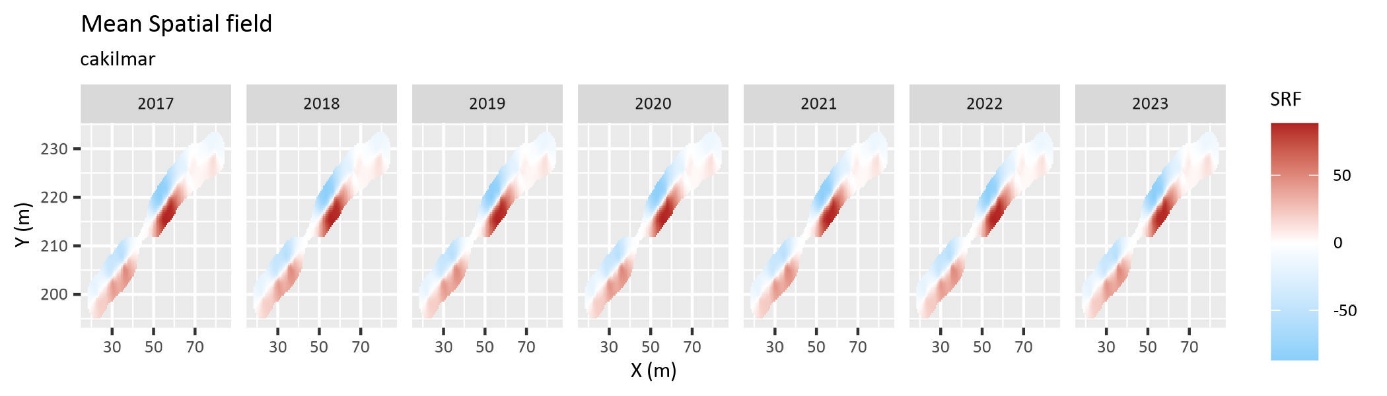
 Fig. S.2.3. Mean spatial field of Cakile maritima occurrence model for each sampling year.

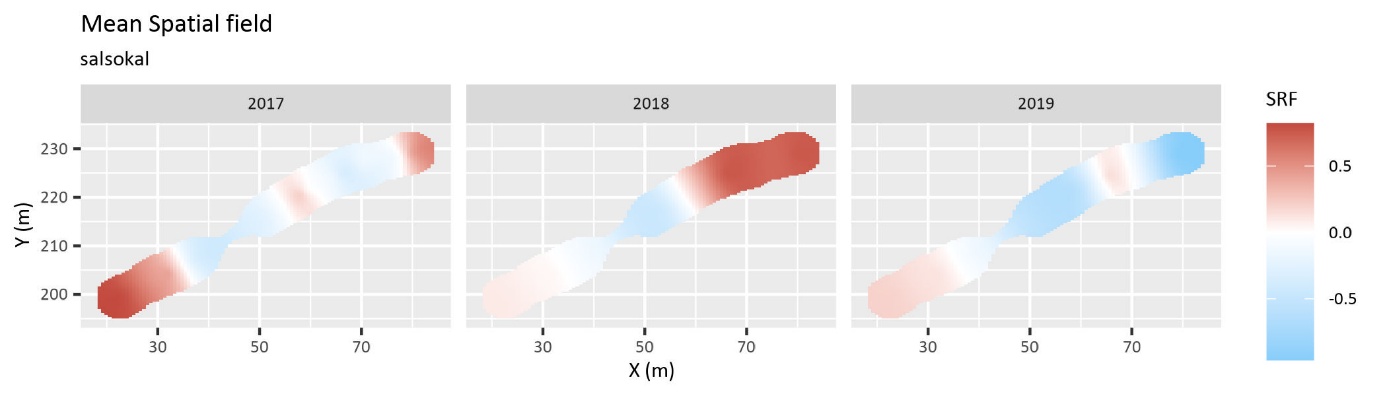
 Fig. S.2.4. Mean spatial field of Salsola kali occurrence model for each sampling year.

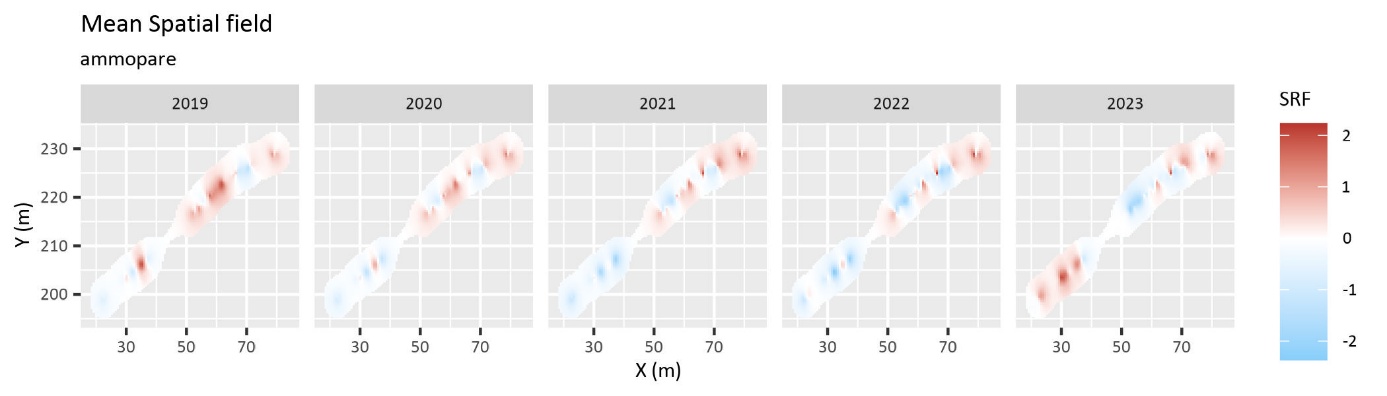
 Fig. S.2.5. Mean spatial field of Calamagrostis arenaria occurrence model for each sampling year.

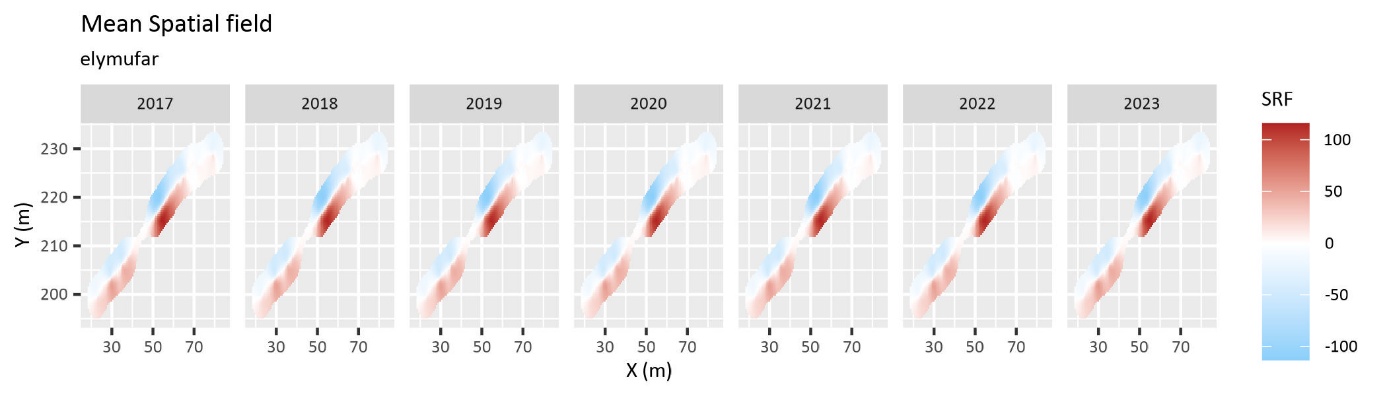
 Fig. S.2.6. Mean spatial field of Elymus farctus occurrence model for each sampling year.

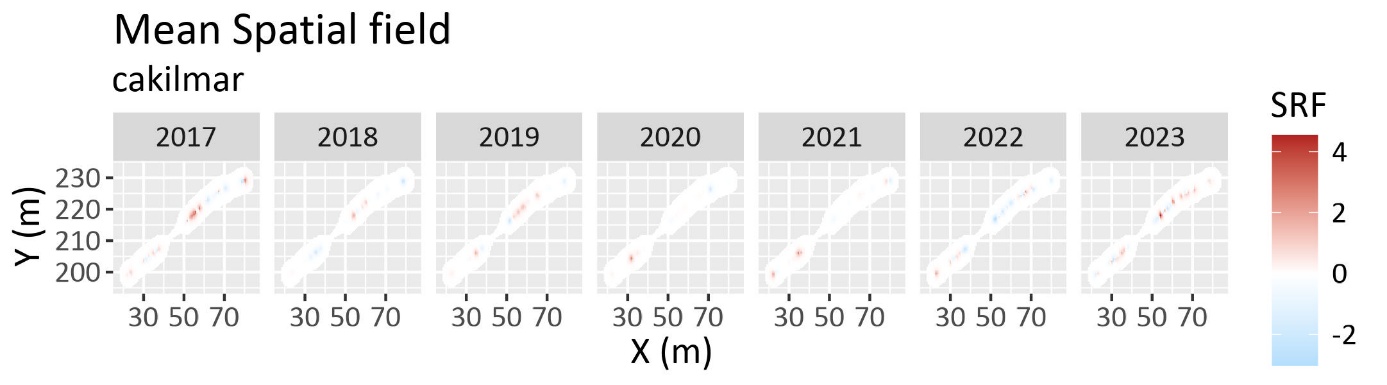
 Fig. S.2.7. Mean spatial field of Cakile maritima abundance/cover model for each sampling year. The spatial field indicates how spatial autocorrelation in the dataset affects the response variable (the plant abundance), in addition to the effect of the covariates which have been accounted for in the model. Positive values increase (overestimate) the response compared to the average, negative values decrease (underestimate) the response.

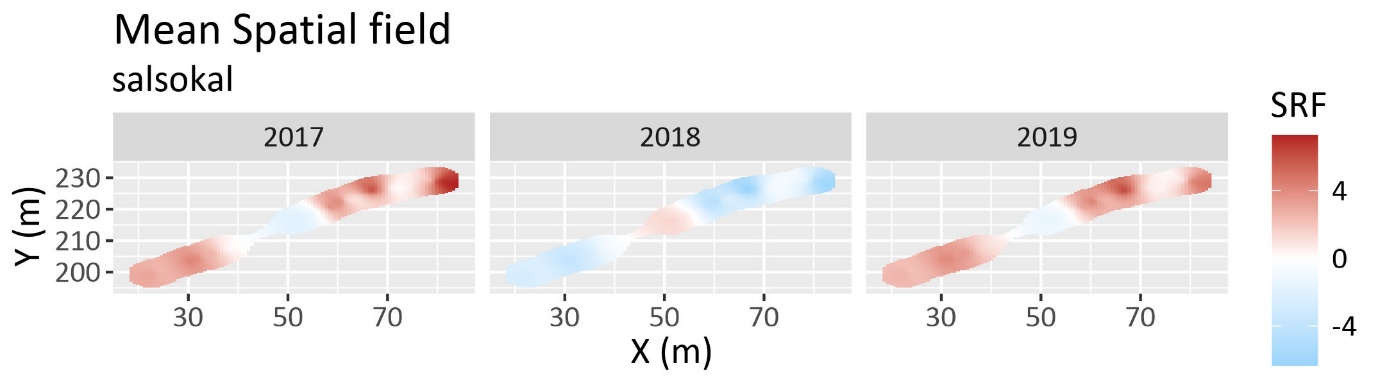
 Fig. S.2.8. Mean spatial field of Salsola kali abundance/cover model for each sampling year.

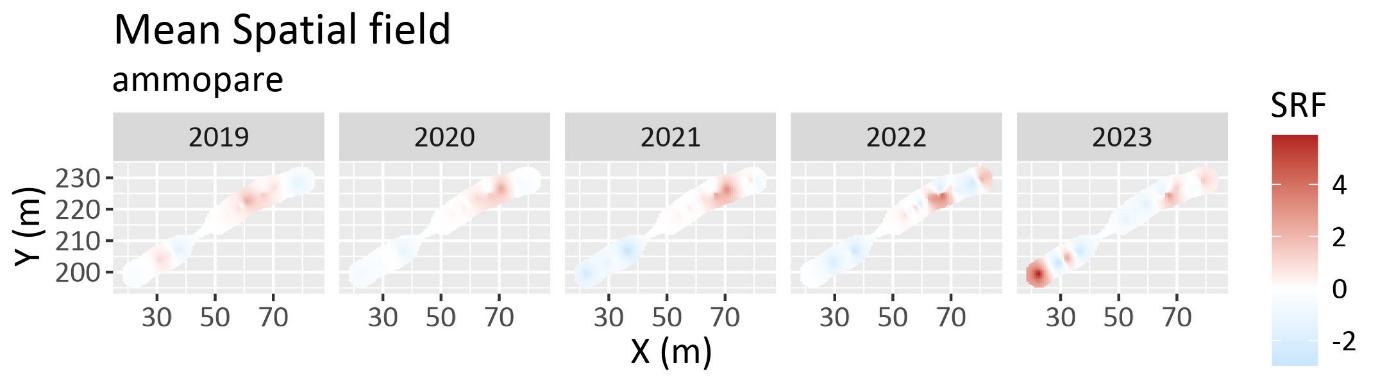
 Fig. S.2.9. Mean spatial field of Calamagrostis arenaria abundance/cover model for each sampling year.

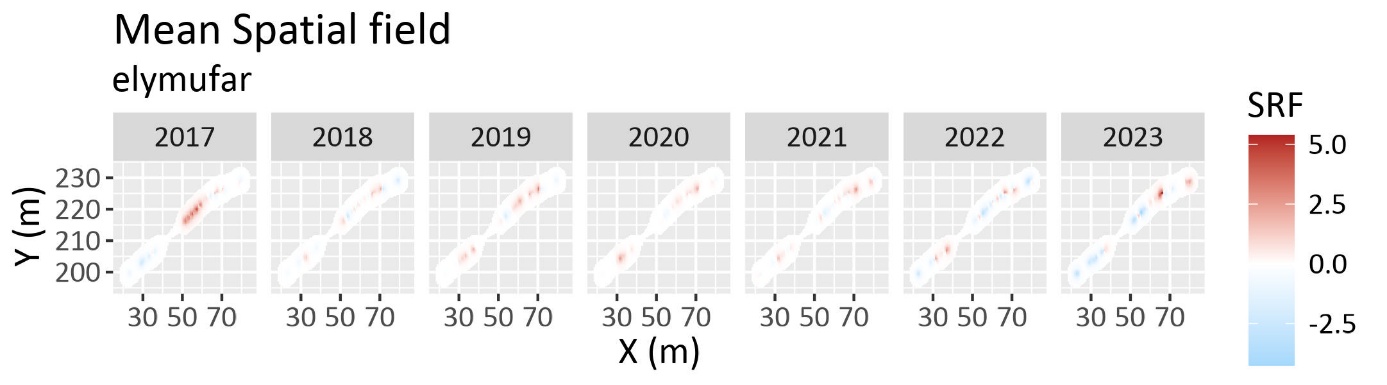
 Fig. S.2.10. Mean spatial field of Elymus farctus abundance/cover model for each sampling year.

### S.3 Spatial predictions

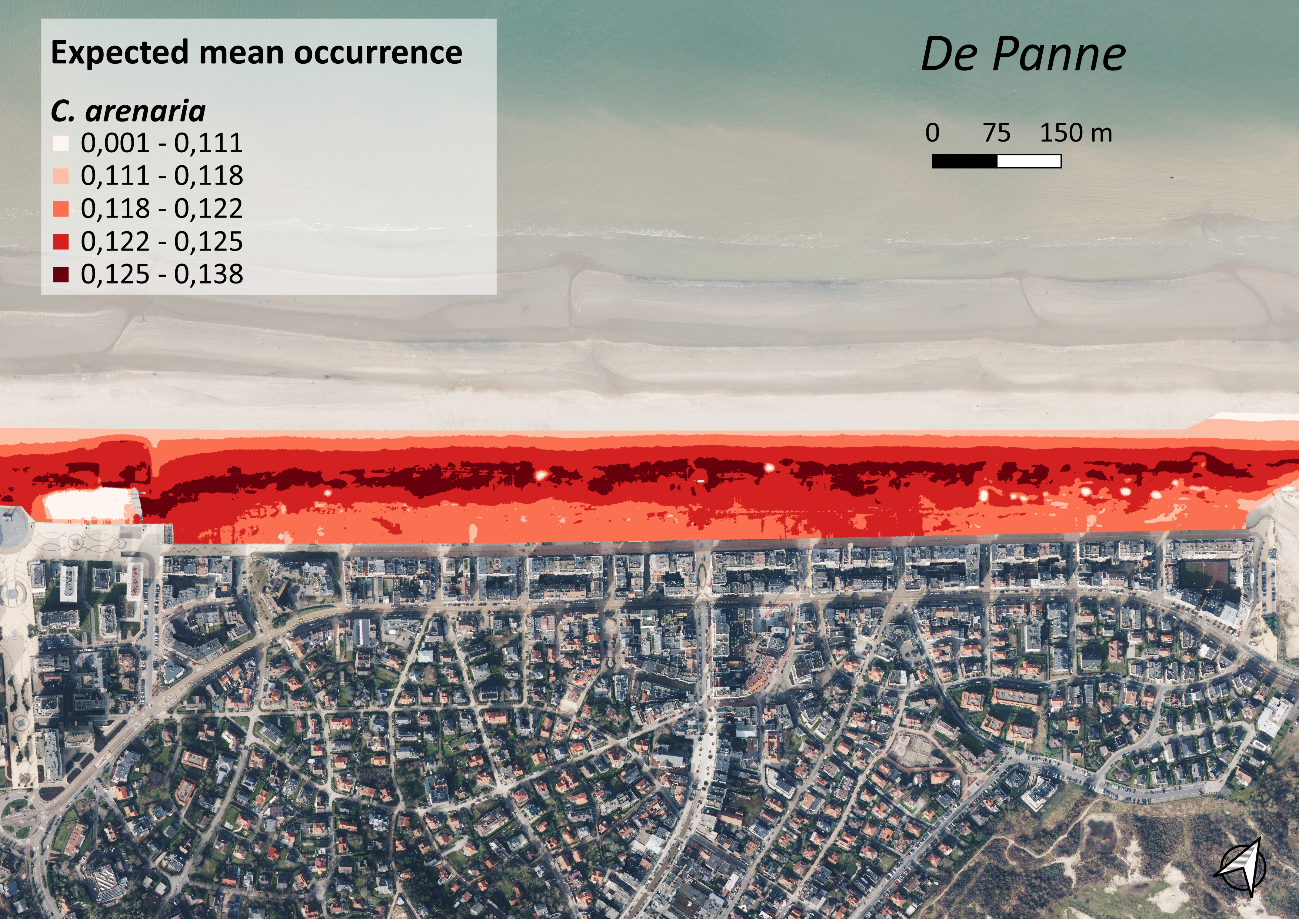

Fig. S.3.1. Expected occurrence probability of C. arenaria on De Panne.

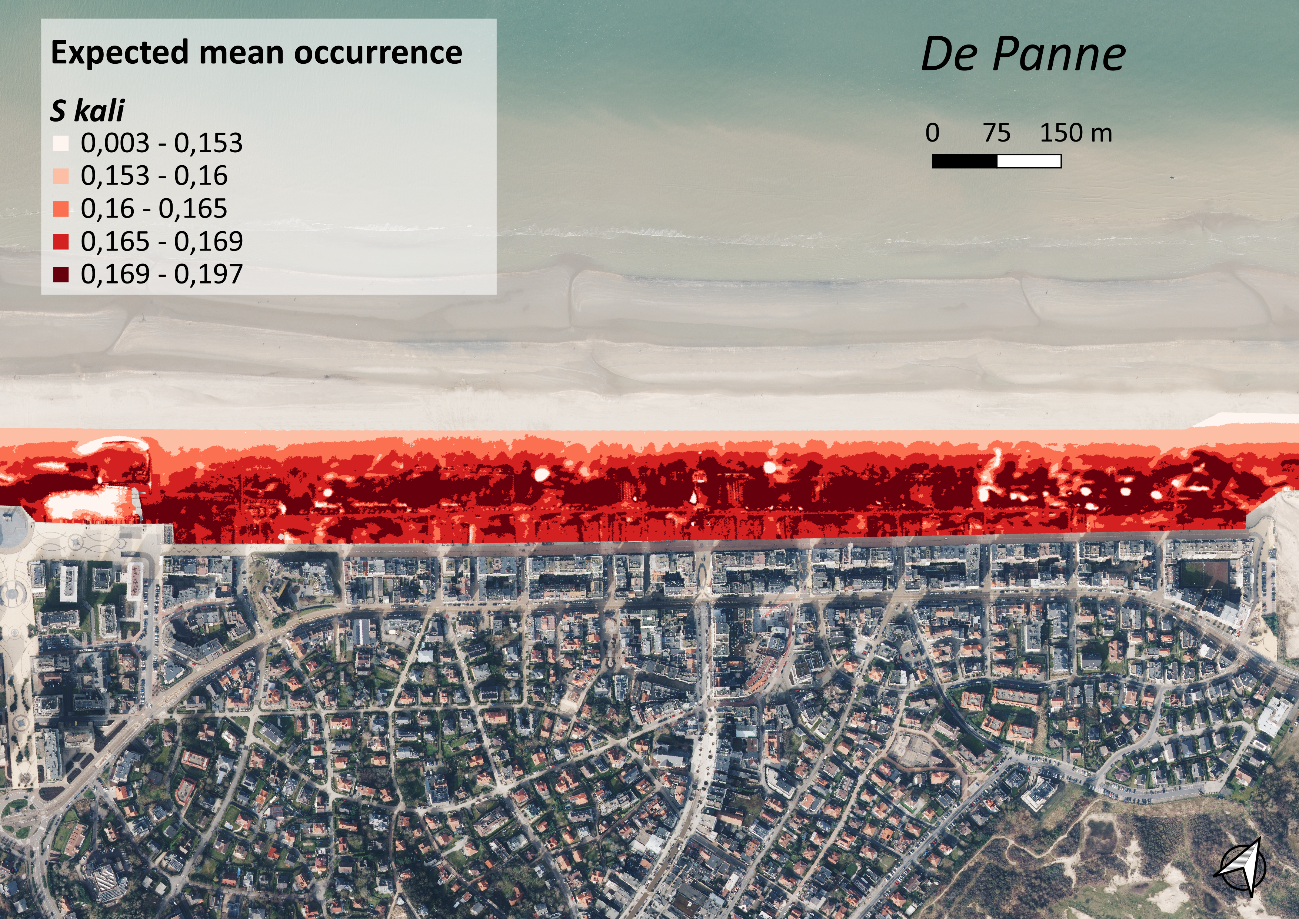

Fig. S.3.2. Expected occurrence probability of S. kali on De Panne.

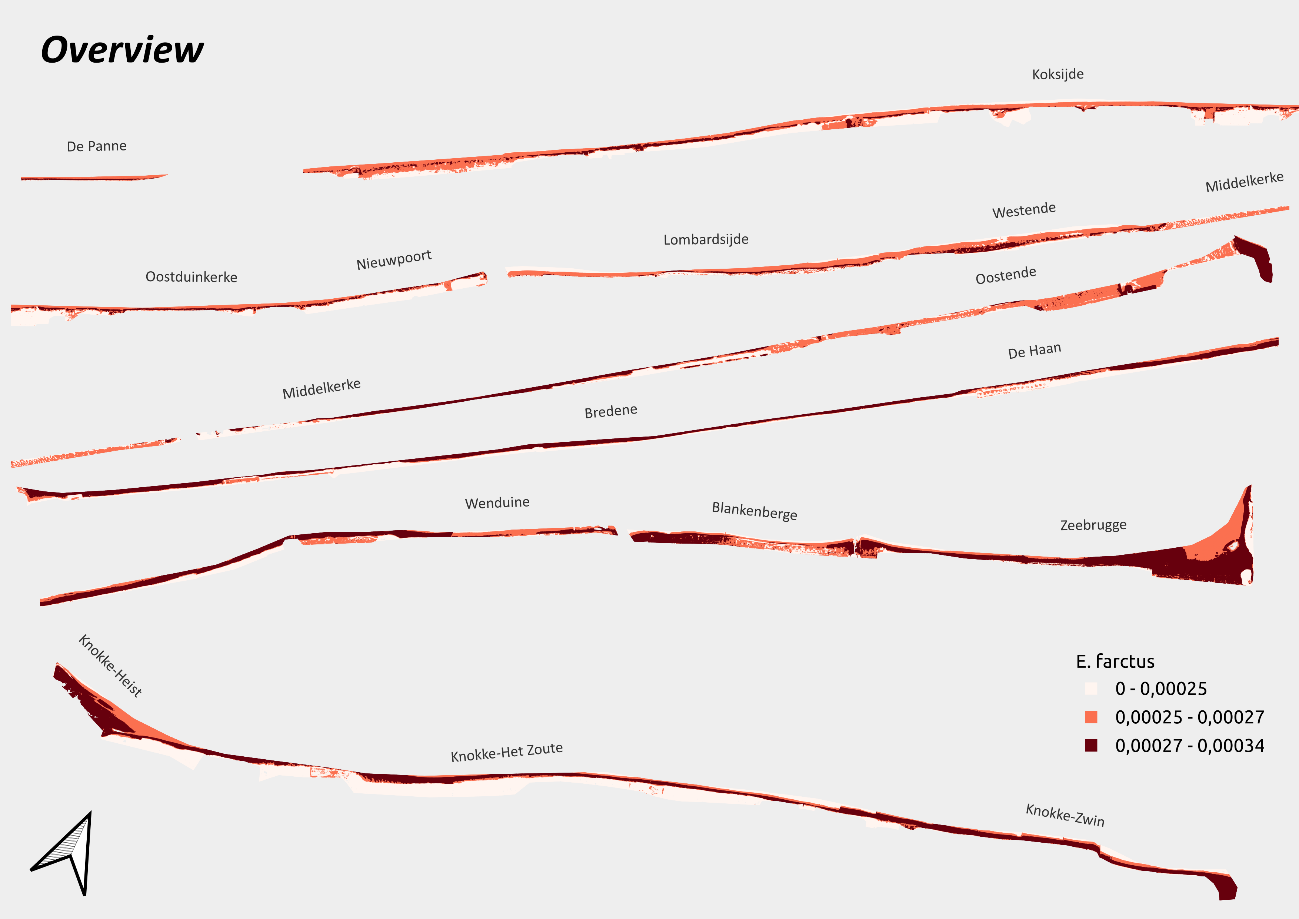

Fig. S.3.3. Expected occurrence probability of E. farctus on the Belgian coastline.

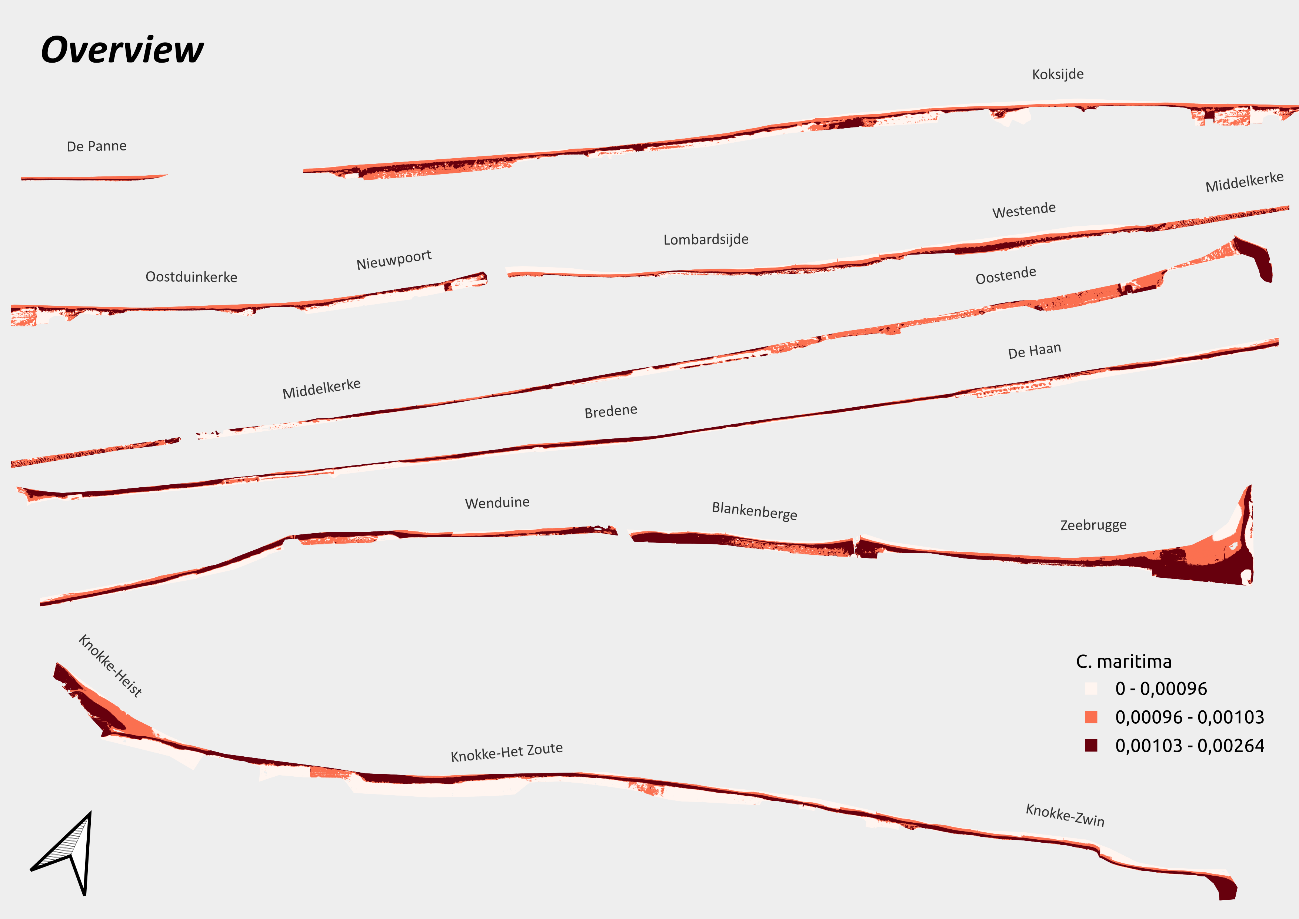

Fig. S.3.4. Expected occurrence probability of C. maritima on the Belgian coastline.

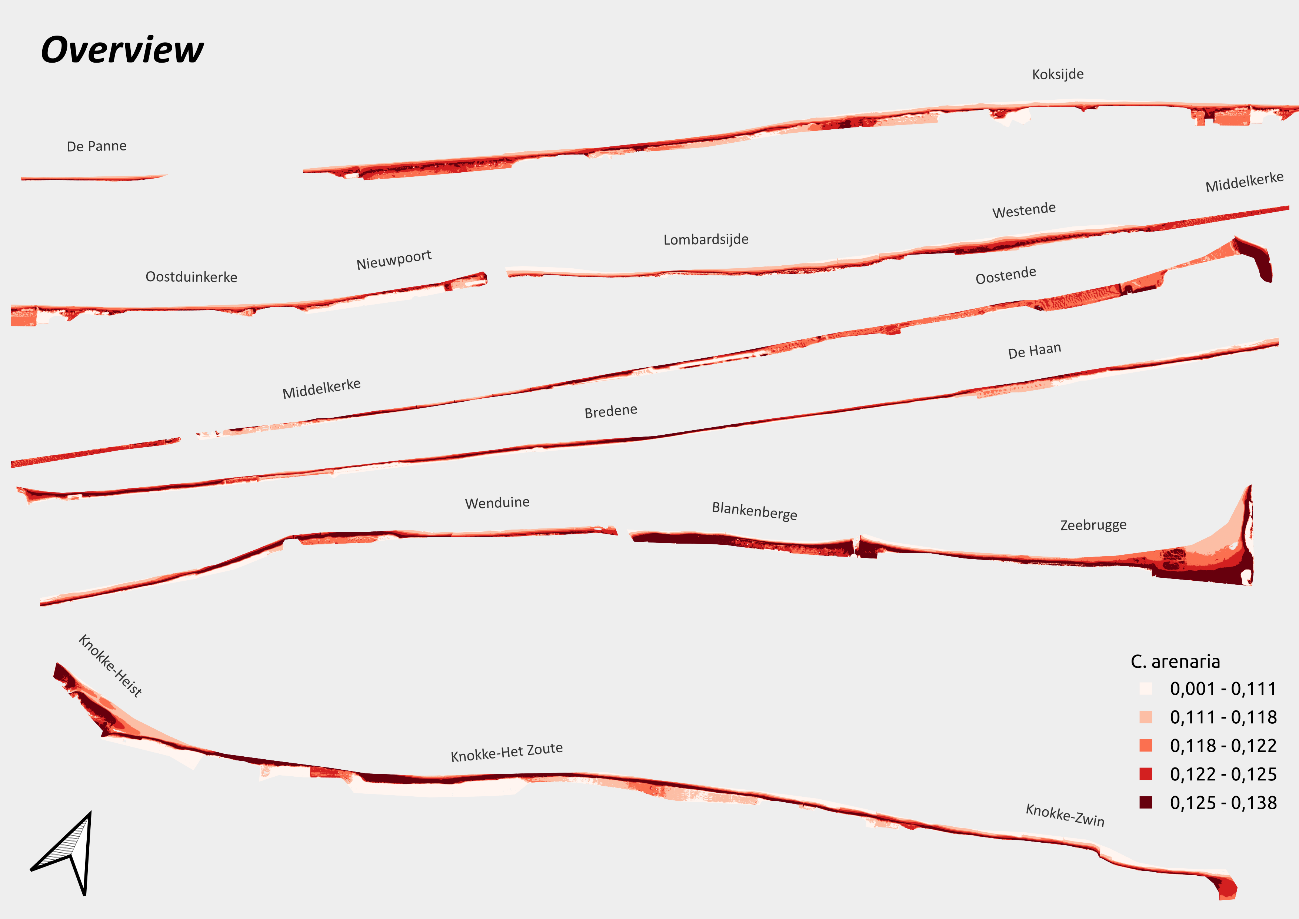

Fig. S.3.5. Expected occurrence probability of C. arenaria on the Belgian coastline.

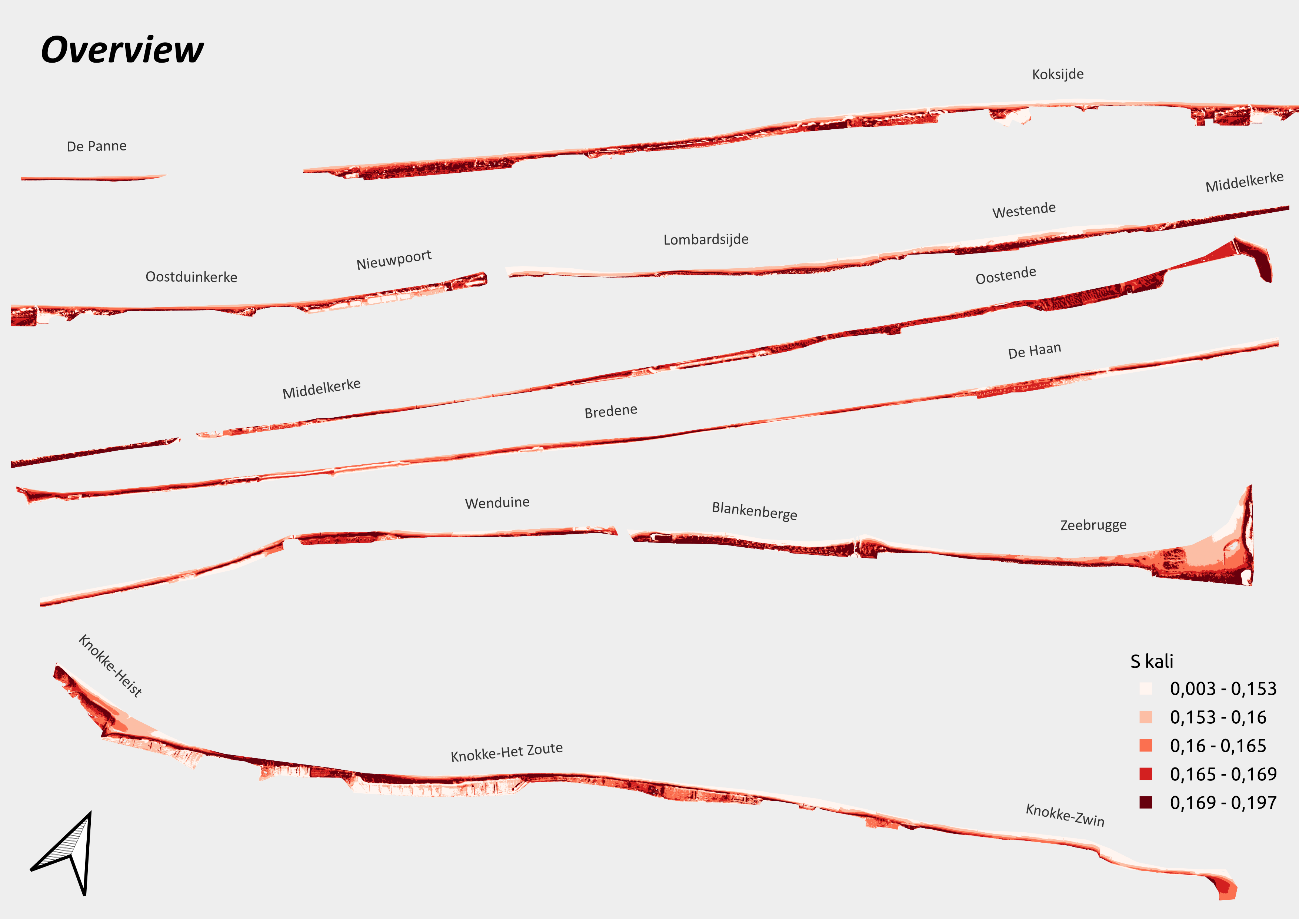

Fig. S.3.6. Expected occurrence probability of S. kali on the Belgian coastline.

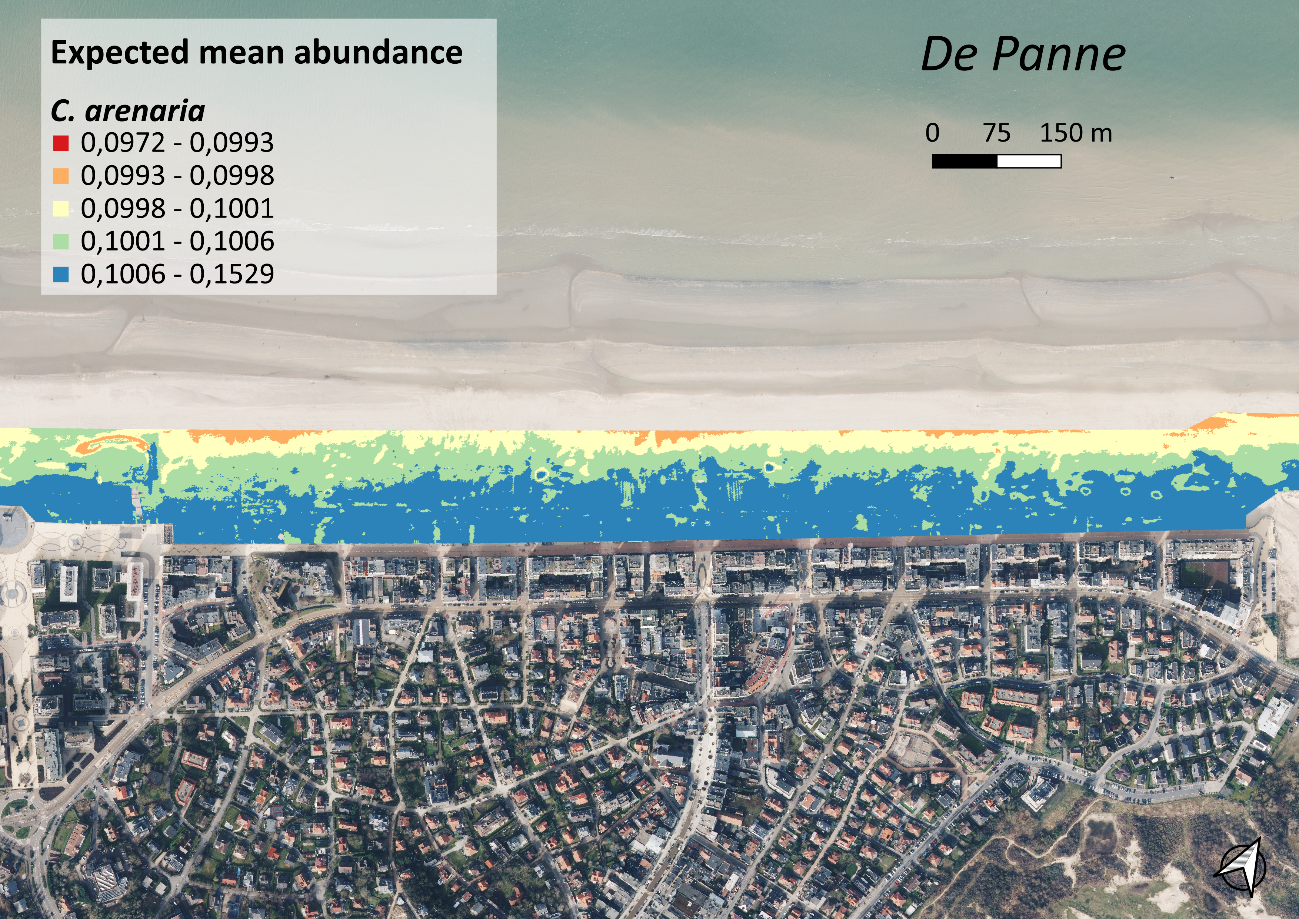

Fig. S.3.7. Expected abundance/cover of C. arenaria on the beach of De Panne.

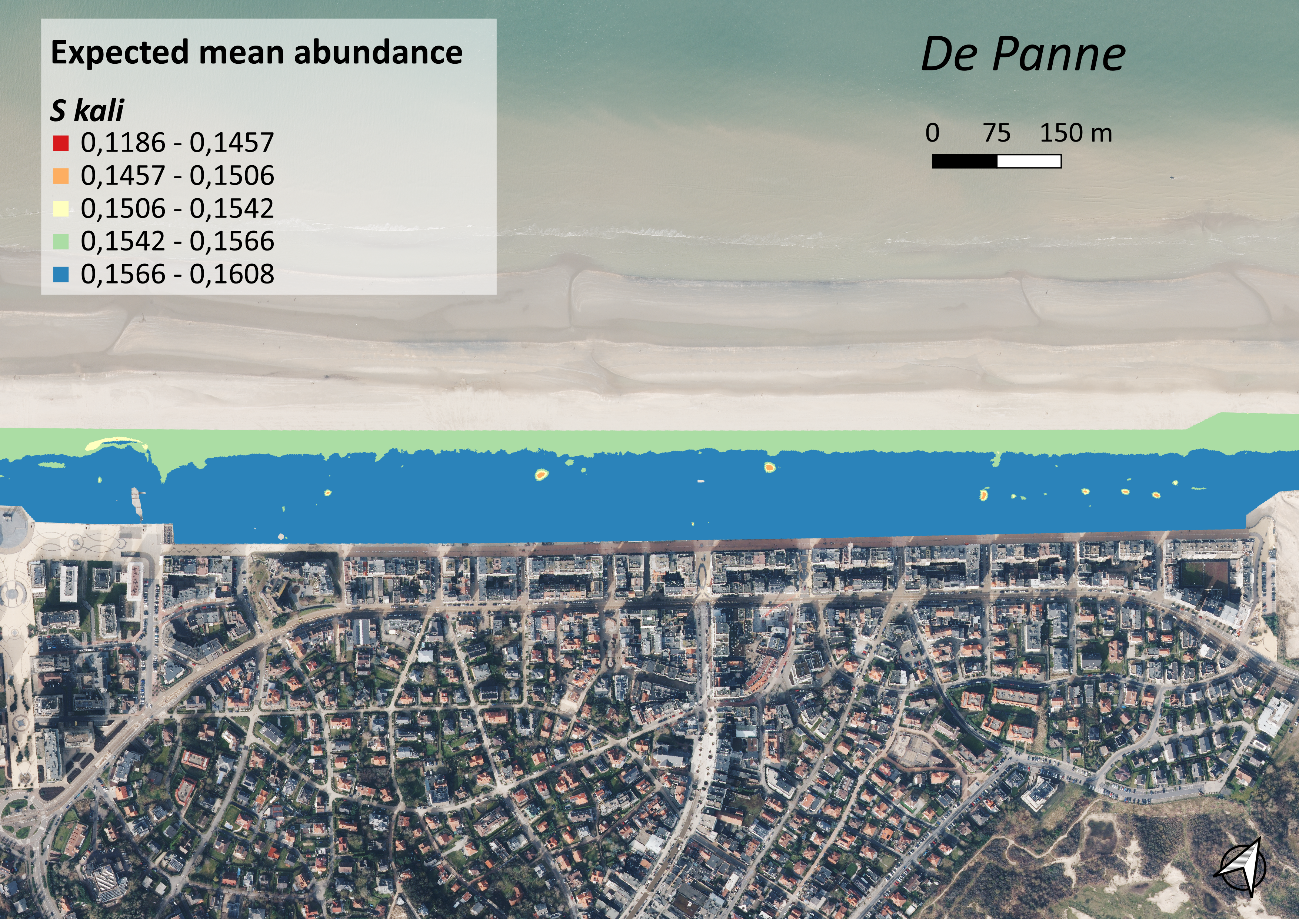

 Fig. S.3.8. Expected abundance/cover of S. kali on the beach of De Panne.

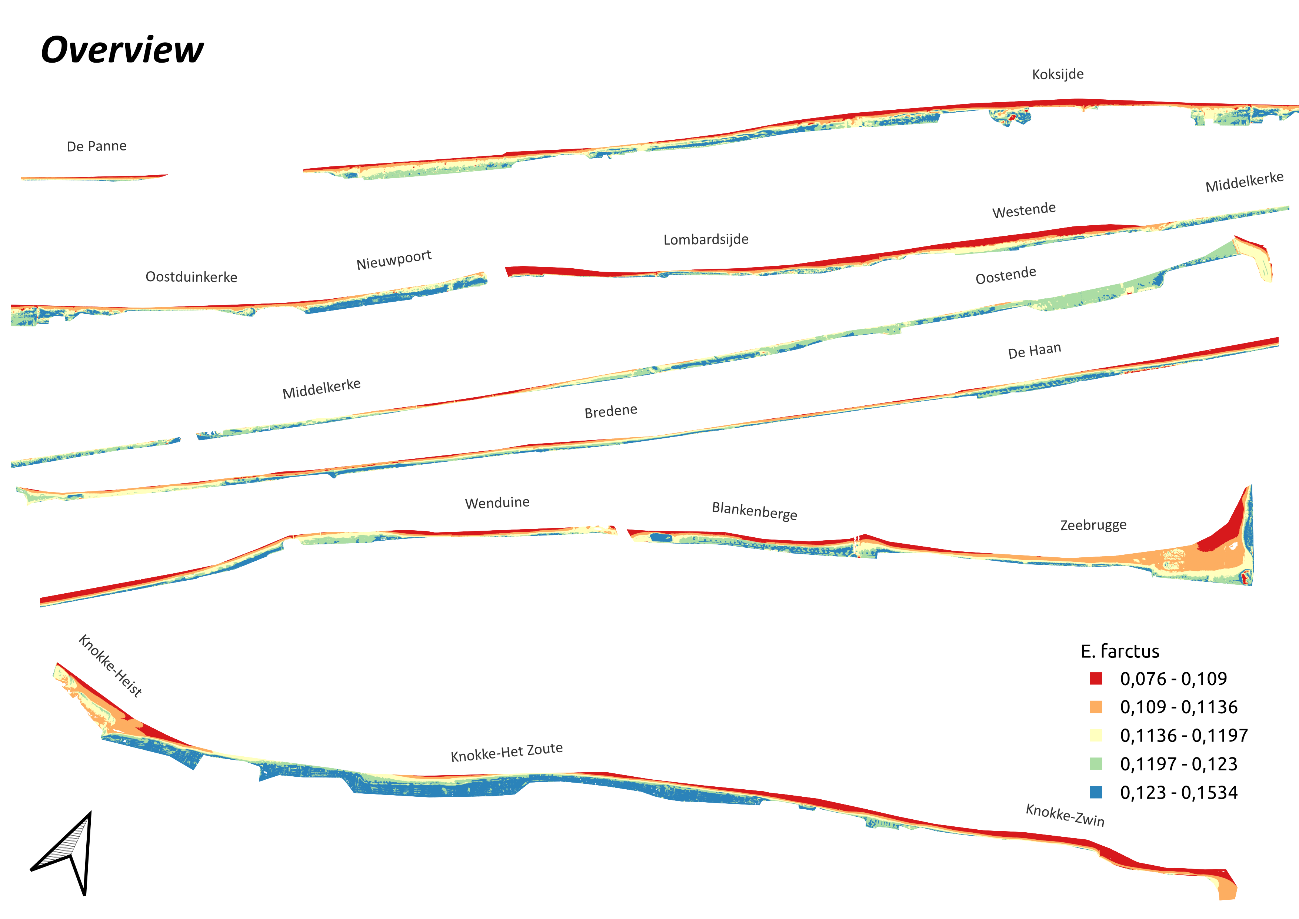

Fig. S.3.9. Expected abundance/cover of E. farctus on the Belgian coastline.

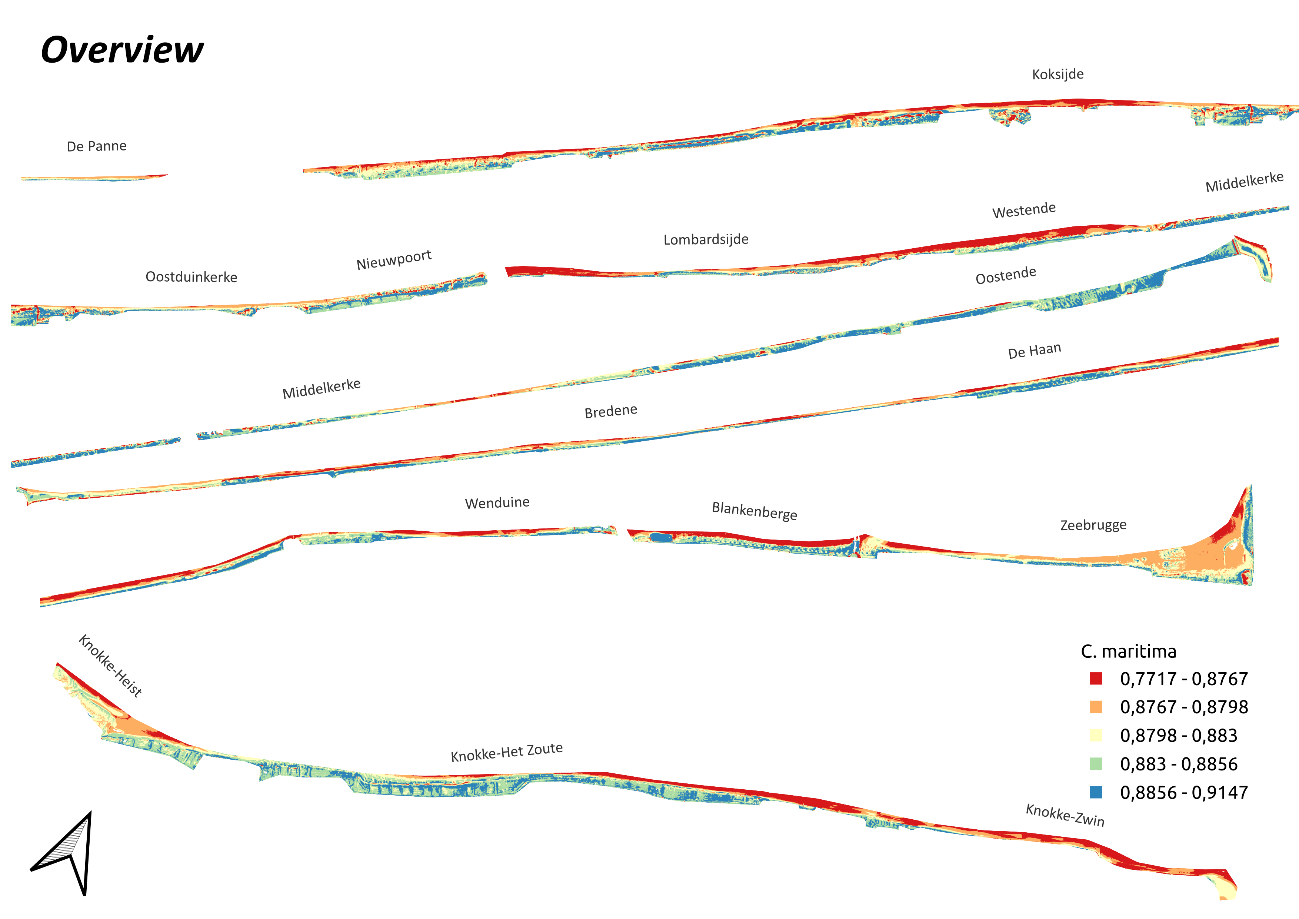

Fig. S.3.10. Expected abundance/cover of C. maritima on the Belgian coastline.
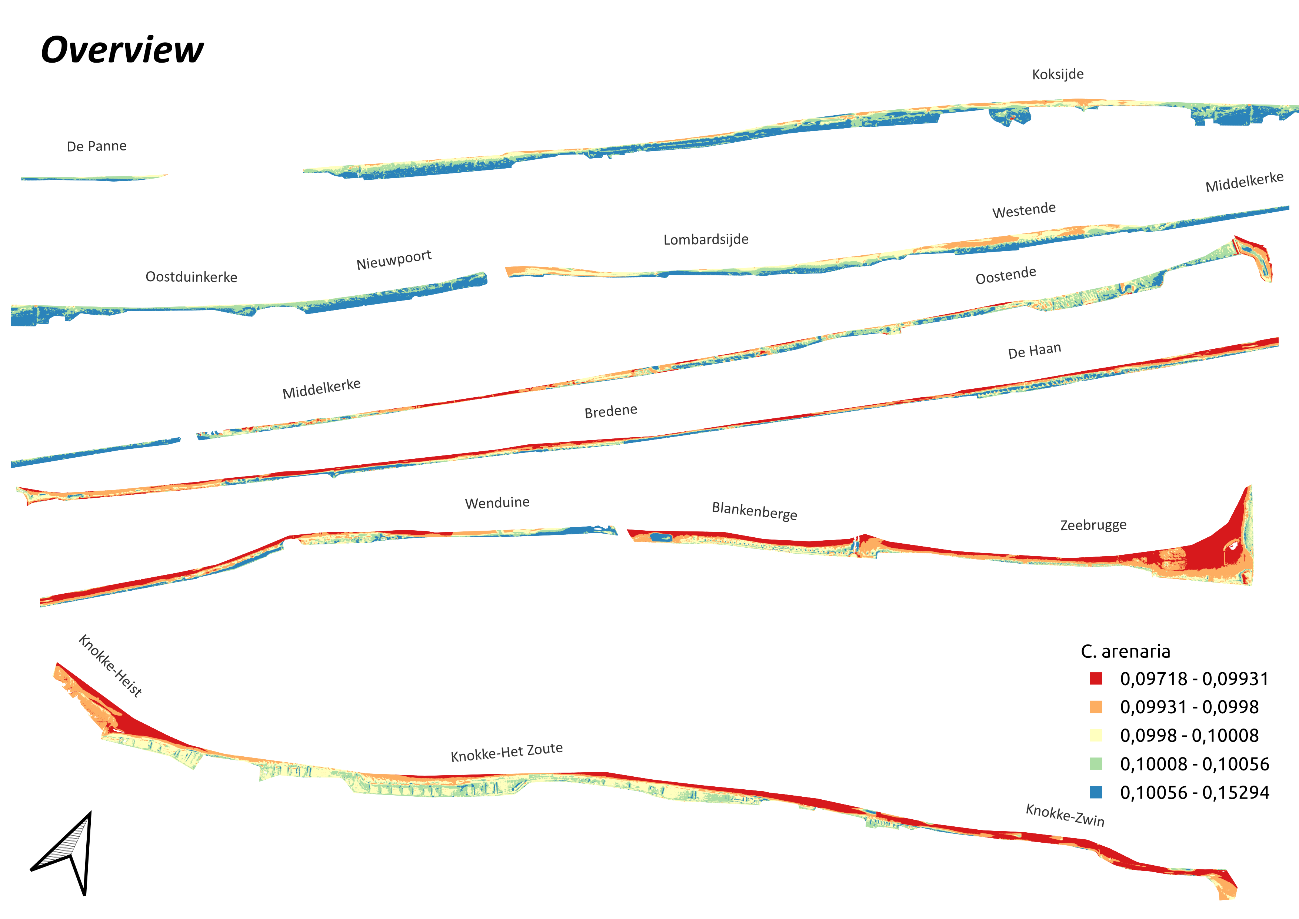

Fig. S.3.11. Expected abundance/cover of C. arenaria on the Belgian coastline.

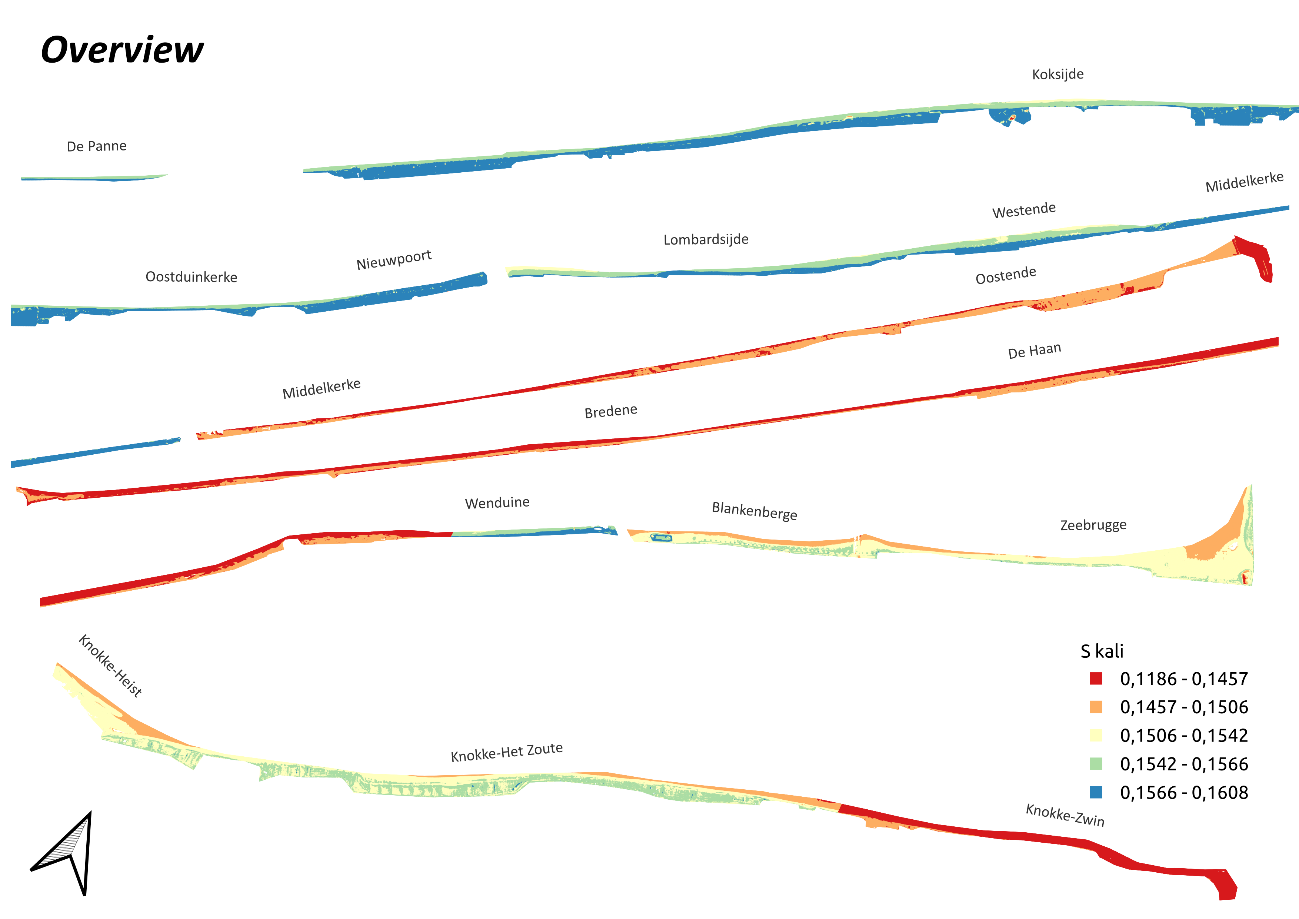

Fig. S.3.12. Expected abundance/cover of S. kali on the Belgian coastline.

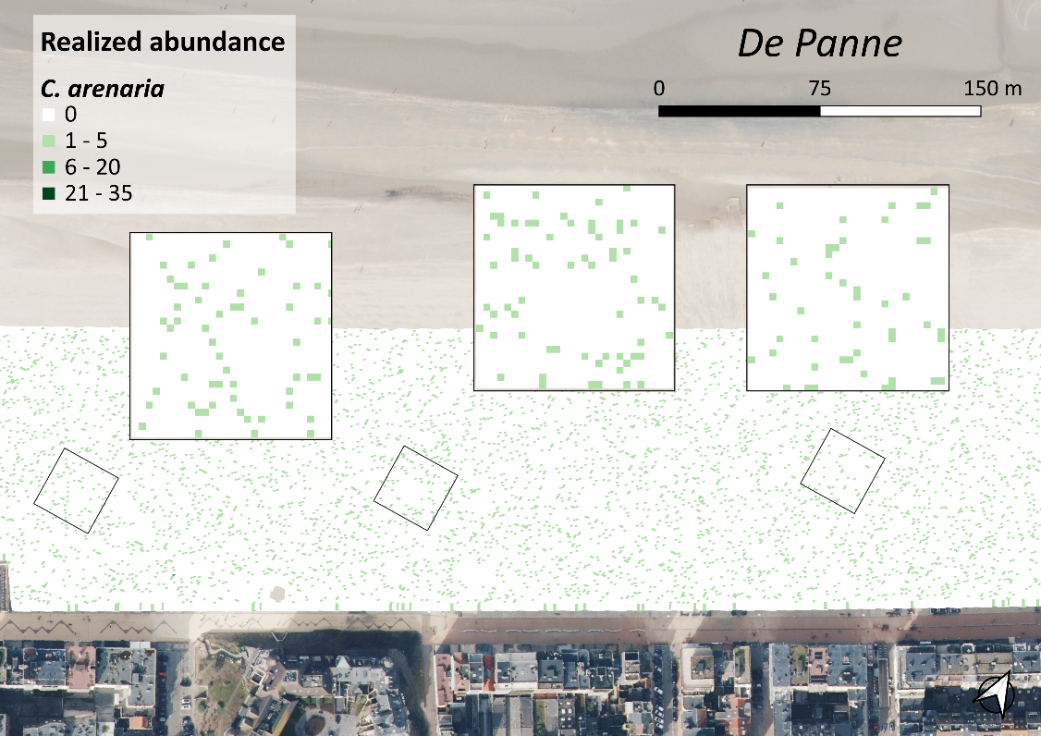

Fig. S.3.13. Simulated abundances/cover of C. arenaria on the beach of De Panne.

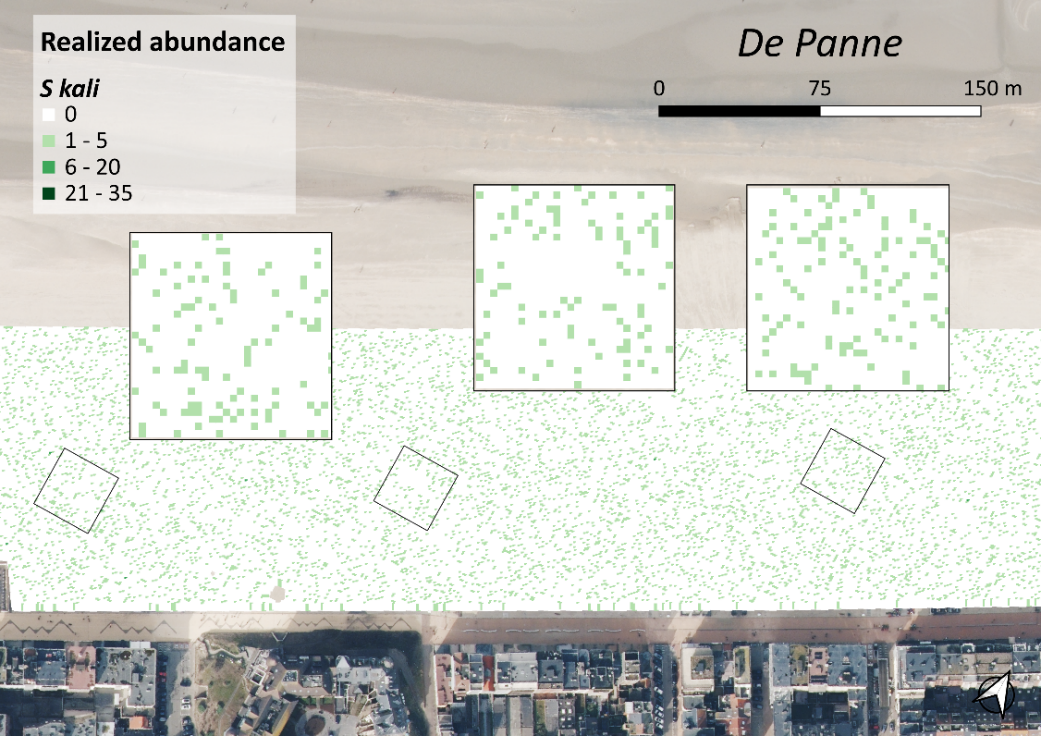

Fig. S.3.14. Simulated abundances/cover of C. arenaria on the beach of De Panne.

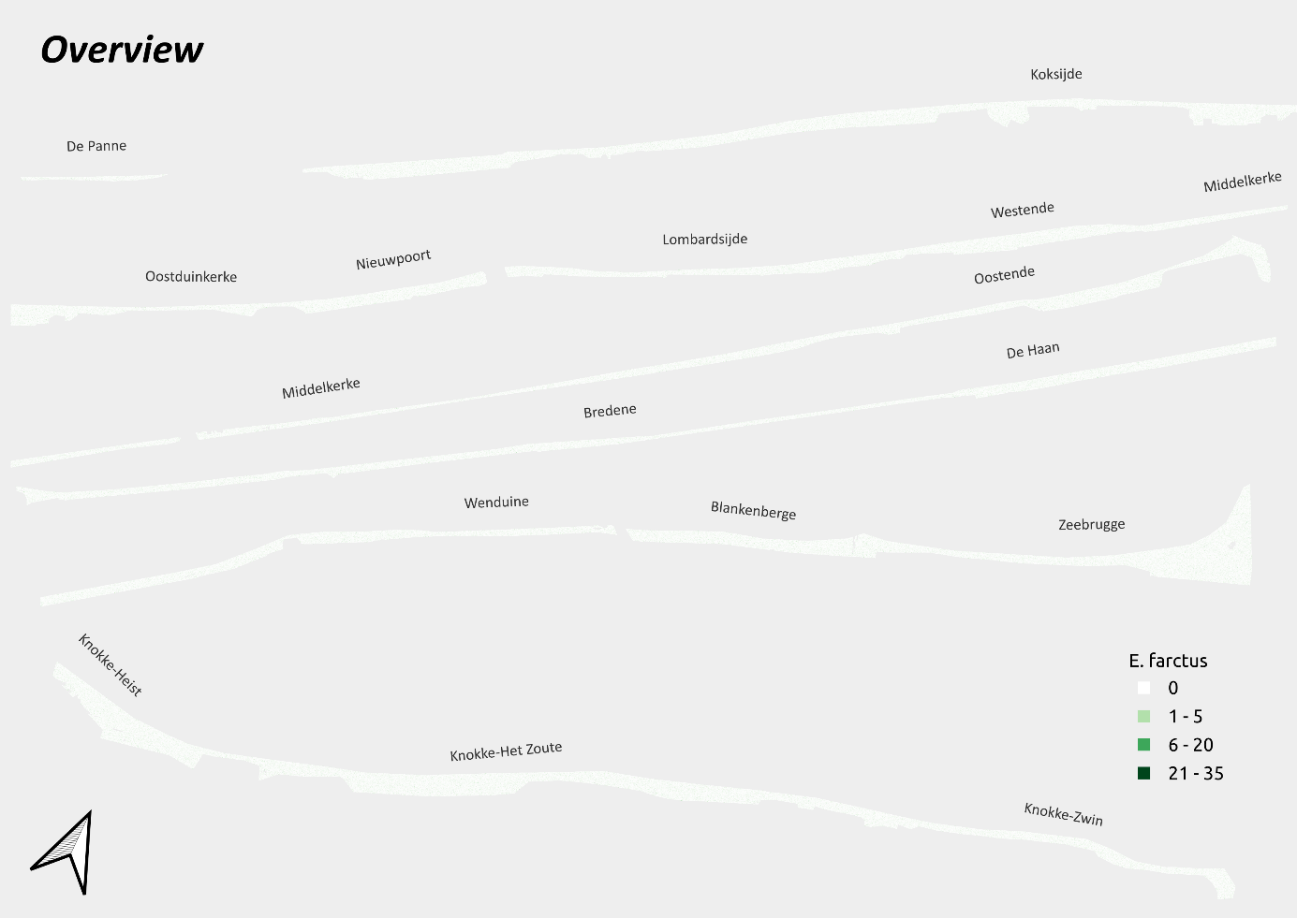

Fig. S.3.15. Simulated abundances/cover of E. farctus on the Belgian coastline.

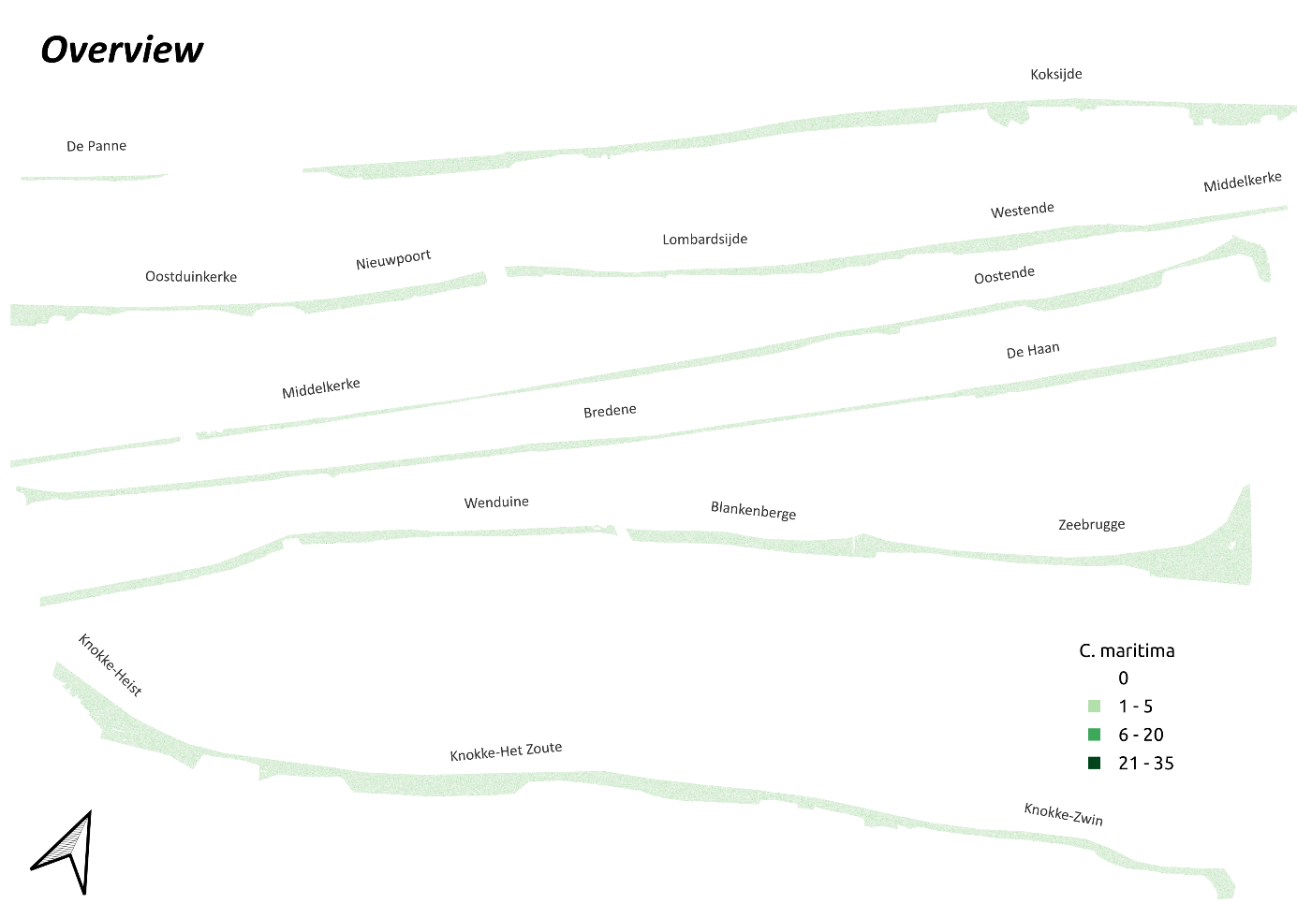

Fig. S.3.16. Simulated abundances/cover of C. maritima on the Belgian coastline.

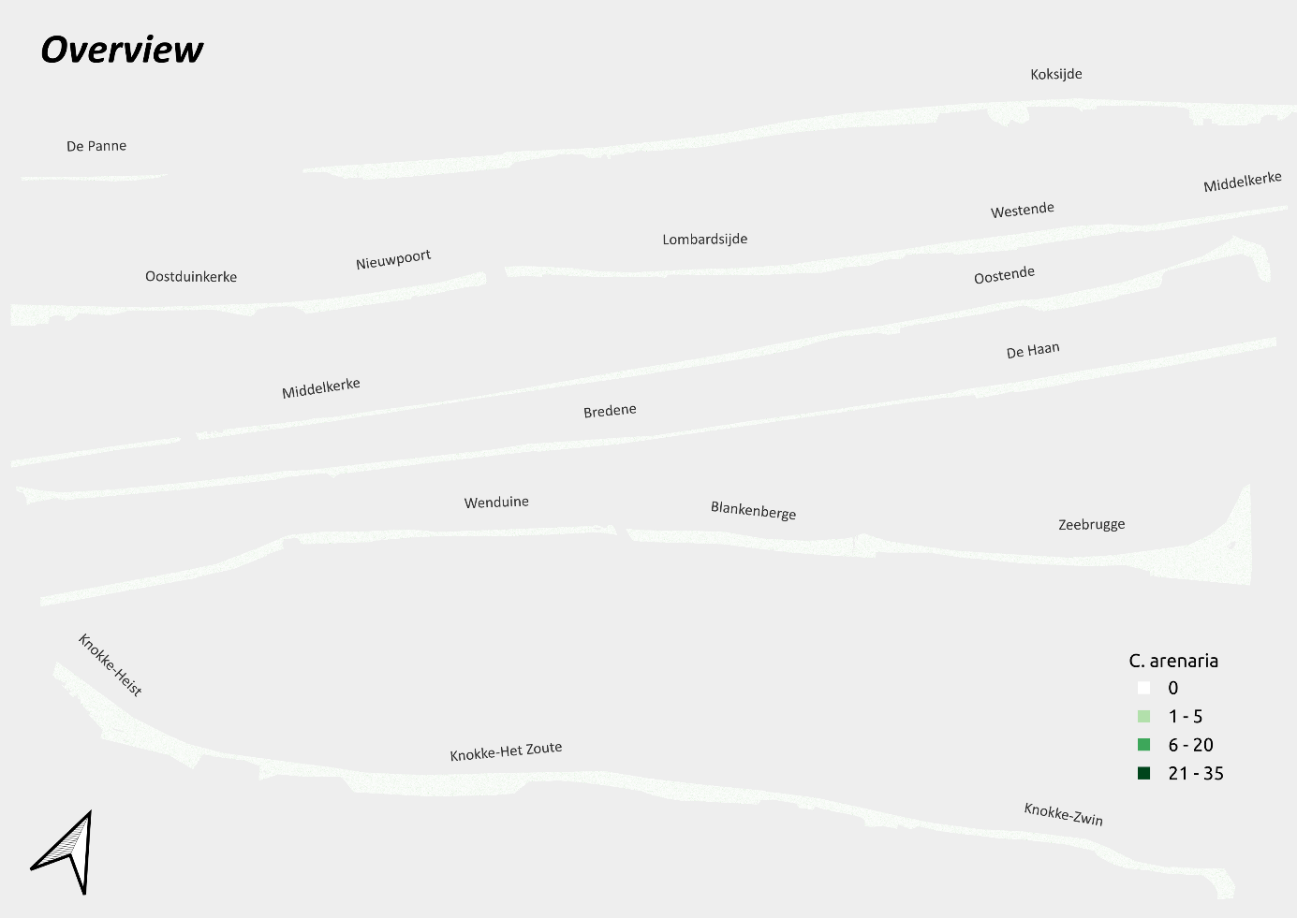

Fig. S.3.17. Simulated abundances/cover of C. arenaria on the Belgian coastline.

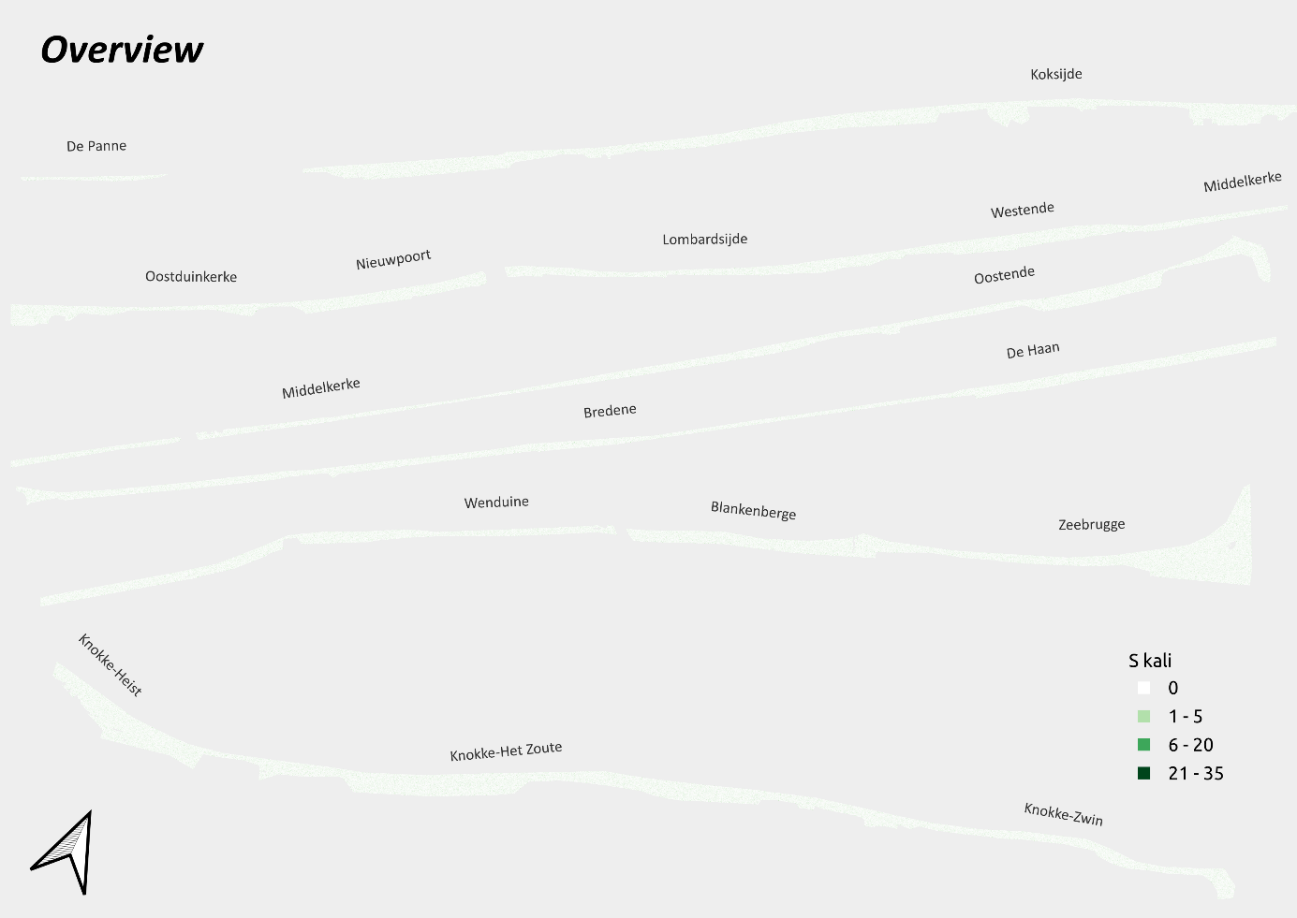

Fig. S.3.18. Simulated abundances/cover of S. kali on the Belgian coastline.

### S.4 Sand accumulation

Table S.4.1. R-output of linear regressions of plant-sand accumulation

**[1] "cakile_maritima"**

 lm(Volume_m3/Surface_m2 ~ 0 + sqrt(Plant_cover_m2),
 data = data)

 Residuals:
 Min	1Q	Median	3Q	Max
 -0.096539 -0.016235 -0.001527 0.023811 0.119062

 Coefficients:
 Estimate Std. Error t value Pr(>|t|)
 sqrt(Plant_cover_m2) 0.158515	0.005786	27.4	<2e-16 ***

 Residual standard error: 0.03617 on 125 degrees of freedom
 Multiple R-squared: 0.8572, Adjusted R-squared: 0.8561
 F-statistic: 750.6 on 1 and 125 DF, p-value: < 2.2e-16
  **[2] "elymus_farctus"**

 lm(Volume_m3/Surface_m2 ~ 0 + sqrt(Plant_cover_m2),
 data = data)

 Residuals:
 Min	1Q	Median	3Q	Max
 -0.091328 -0.021595 -0.006558 0.009706 0.161484

 Coefficients:
 Estimate Std. Error t value Pr(>|t|)
 sqrt(Plant_cover_m2) 0.15542	0.00642	24.21	<2e-16 ***

Residual standard error: 0.03841 on 89 degrees of freedom
 Multiple R-squared: 0.8682, Adjusted R-squared: 0.8667
 F-statistic: 586.1 on 1 and 89 DF, p-value: < 2.2e-16

 **[3] "calamagrostis_arenaria"**

 lm(Volume_m3/Surface_m2 ~ 0 + sqrt(Plant_cover_m2),
 data = data)

 Residuals:
 Min	1Q	Median	3Q	Max
 -0.016018 -0.005992 -0.001895 0.004725 0.017892

 Coefficients:
 Estimate Std. Error t value Pr(>|t|)
 sqrt(Plant_cover_m2) 0.14221	0.00953	14.92 1.46e-06 ***

 Residual standard error: 0.01109 on 7 degrees of freedom
 Multiple R-squared: 0.9695, Adjusted R-squared: 0.9652
 F-statistic: 222.7 on 1 and 7 DF, p-value: 1.455e-06

 **[4] "salsola_kali"**

 lm(Volume_m3/Surface_m2 ~ 0 + sqrt(Plant_cover_m2),
 data = data)

 Residuals:
 Min	1Q	Median	3Q	Max
 -0.078915 -0.006934 0.005945 0.016062 0.091890

 Coefficients:
 Estimate Std. Error t value Pr(>|t|)
 sqrt(Plant_cover_m2) 0.12076	0.01393	8.671 1.63e-06 ***

 Residual standard error: 0.04426 on 12 degrees of freedom
 Multiple R-squared: 0.8624, Adjusted R-squared: 0.8509
 F-statistic: 75.19 on 1 and 12 DF, p-value: 1.633e-06
